# Quantifying Functional Conservation of Human and Mouse Regulatory Elements via FUNCODE

**DOI:** 10.1101/2024.10.31.620766

**Authors:** Weixiang Fang, Chaoran Chen, Boyang Zhang, Yi Wang, Ruzhang Zhao, Weiqiang Zhou, Hongkai Ji

**Author notes:** Correspondence (H.J.).

## Abstract

Evolutionary conservation is crucial for understanding genome functions and lays the foundation for using animal models in studying human diseases. However, conventional conservation scores based on DNA sequence evolution do not capture the dynamic biochemical activities of DNA elements, termed functional conservation. Quantifying functional conservation has been limited by the availability of functional genomic data matched across species. To address this, we developed FUNCODE, a framework for characterizing functional conservation through *in silico* sample matching. Applying FUNCODE to 2,595 uniformly processed datasets from the Encyclopedia of DNA Elements (ENCODE), we generated genome-wide FUNCODE scores for human and mouse regulatory elements, identifying 3.3 million functionally conserved human-mouse element pairs. We demonstrate FUNCODE’s diverse applications, including annotating 78,501 novel regulatory elements, transferring 37,968 high-resolution human ENCODE Hi-C loops in immune lineages to mice, identifying conserved functional signals for disease modeling, and enhancing cross-species integration of single-cell omics data.

## Introduction

Comparative genomics, the practice of comparing genome sequences across different species, has been instrumental in studying evolution, identifying functional DNA elements in genomes, and building animal models of human diseases^1,2^. Building upon the principle that negative (purifying) selection on DNA elements with important functions results in conserved DNA sequences across species, sequence conservation scores, such as PhastCons^3^ and PhyloP^4^, have been essential for facilitating these applications. However, it is known that DNA elements with conserved sequences can have diverging functions, evidenced by differences in tissue or cell-type-specific chromatin accessibility^5,6^, histone modifications^7,8^, and transcription factor binding profiles^9,10^. Specifically, regulatory elements can acquire new functions, be repurposed within large combinatorial networks or modules, or change their roles depending on the genomic contexts in different species^11–14^. As a result, cross-species analysis based solely on sequence conservation can lead to inaccurate conclusions when the functional outcome is concerned. Therefore, to better characterize functional conservation, defined here as conservation in DNA elements’ dynamic biochemical activities, and transfer knowledge between animal models and human studies, there is a need for genome-wide conservation scores grounded on functional genomics data.

The challenge in studying the functional conservation of regulatory elements lies in the dynamic nature of their activities. When researchers use an animal model to validate a human regulatory element, they strive to replicate relevant factors such as tissue type, cell type, developmental stage, and environment, operating under the hypothesis that the regulatory element’s function is conserved. The objective is to test and reproduce the element’s activity under these controlled conditions. Consequently, defining global functional conservation necessitates a holistic consideration of context across the entire organism and over an extended timescale.

Despite the extensive generation of functional genomics data over the years, cross-species comparative analyses have largely been restricted to a limited subset of tissue or cell types, often paired manually. For example, the original mouse Encyclopedia of DNA Elements (mouseENCODE) compared transcription factor binding, histone modification, and methylation profiles of three paired human and mouse cell lines^10^. Later, the regulatory landscapes of DNase hypersensitivity sites (DHSs) were compared between human and mouse genomes across sixteen tissues, cell lines, or cell types^15^. Similarly, regulatory element activities were examined in five tissues across ten mammalian species^8,16^. However, none of these studies provided a practical conservation score, likely due to the limited number of paired samples involved.

Efforts to develop conservation scores based on functional genomics data have typically focused on predicting the presence or absence of genomics signals, without considering whether the activities are consistent across contexts. This approach can mistakenly label evolutionary repurposed elements – those active in different contexts in different species – as conserved. As a result, these elements are unlikely to reproduce their original human activity in an animal model. For instance, LECIF recently scored functional conservation between human and mouse by predicting whether their elements could be sequence-aligned^17^. However, earlier research estimated that 37% of sequence-aligned human and mouse DHSs were functionally repurposed^15^. Similarly, DeepGCF used a comparable approach to score conservation between humans and pigs^18^.

The Encyclopedia of DNA Elements (ENCODE) consortium, with its extensive collection of functional genomic data for human and mouse, offers a unique opportunity to overcome these limitations^19^ [ENCODE tracking# ENC4P01]. As part of the consortium effort, we developed FUNCODE, a scalable computational framework to score cross-species functional conservation of DNA elements. FUNCODE leverages advancements in integrative genomics to generate numerous *in silico* sample pairs across species, eliminating the need for manual pairing. By applying FUNCODE to ENCODE’s DNase-seq, ATAC-seq, and histone ChIP-seq data, we have produced genome-wide functional conservation scores for human and mouse regulatory elements, providing a foundational tool and resource for cross-species data integration, knowledge transfer, and functional annotation.

## Results

### FUNCODE framework for scoring functional conservation of DNA elements

FUNCODE leverages a compendium of functional genomic data from two species to quantify functional conservation of their DNA elements paired through either sequence homology or gene homology (**Fig. 1a**). We distinguish between two types of conservation. The first, COnservation of Variable Activity (CO-V), describes the conservation of tissue- or cell-type-specific activities. The second, COnservation of Baseline Activity (CO-B), describes conserved constitutive activities (**Fig. 1b**).

**Fig. 1.**
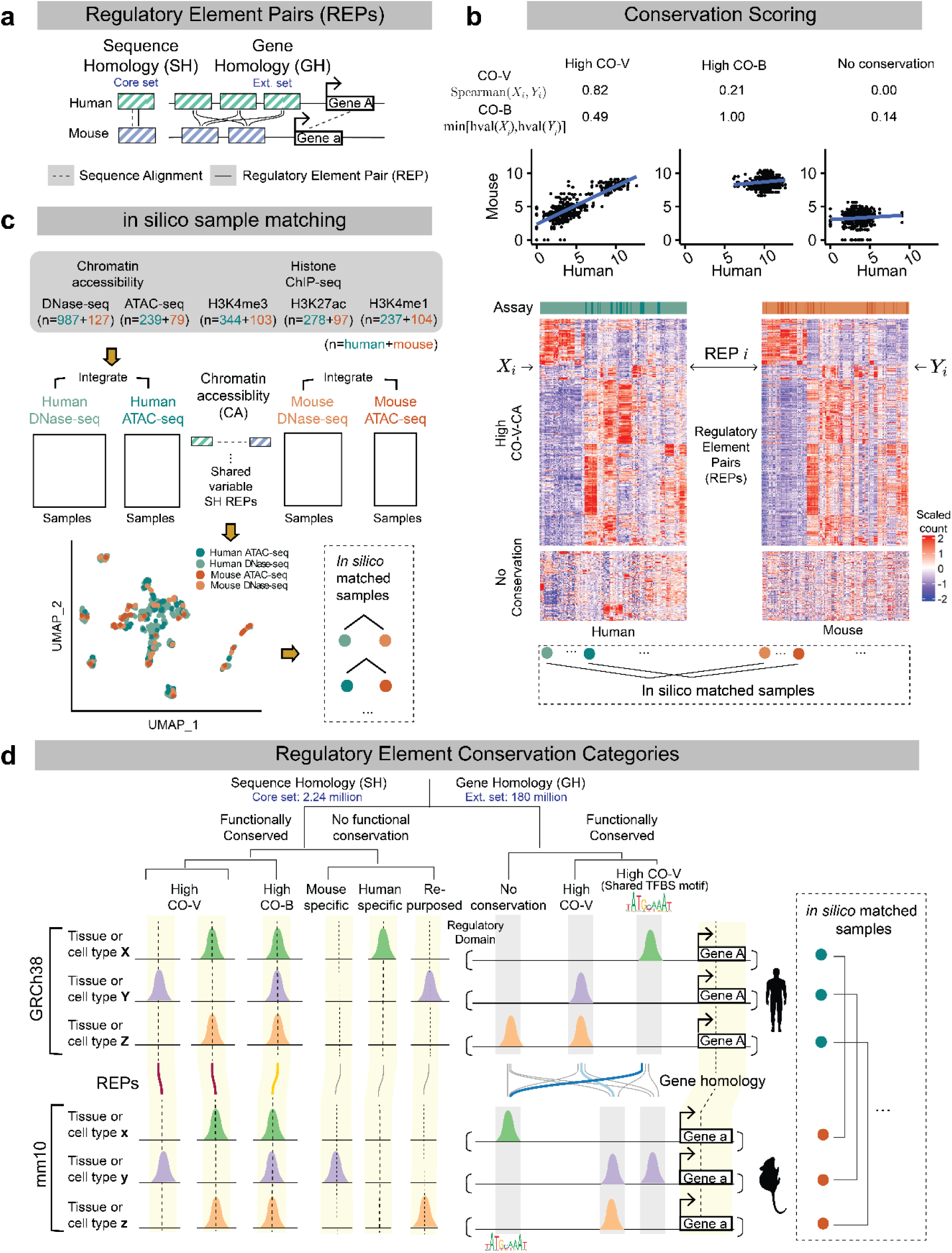
Overview of the FUNCODE framework. (**a**) Schematic illustrating two sets of regulatory element pairs (REPs) distinguished by sequence homology and gene homology. A DNA element is classified in the core set (left) if it can be mapped to another species via sequence alignment, indicating sequence homology. Conversely, a DNA element belongs to the extended set (right) if it pairs with an element near orthologous genes in another species, even without sequence homology, indicating gene homology. (**b**) Schematic illustrating the concepts of Conservation of Variable Activities (CO-V) and Conservation of Baseline Activity (CO-B). Scatterplots demonstrate different conservation patterns, each representing one REP, with data points for paired human and mouse samples. A REP with a high CO-V score shows highly correlated activities across *in silico* matched sample pairs. In contrast, a REP with a high CO-B score exhibits consistently high activity across all tissue or cell types but may have low correlation. REPs with low CO-V and CO-B scores lack conservation. Example heatmaps display REPs (rows) with either high CO-V scores for chromatin accessibility (top) or no conservation (bottom), with columns representing *in silico* matched samples. (**c**) Schematic illustrating the steps of *in silico* sample matching. REPs with sequence homology and highly variable activities across samples in both species were identified and used as features. Functional genomics data were co-embedded into a common low-dimensional space (shown in a 2D scatterplot). In this space, *in silico* matched samples were identified, enabling the calculation of CO-V scores. (**d**) Schematic illustrating the categorization of DNA elements based on their associated REPs and conservation scores. DNA elements linked to core set REPs can have either high CO-V or high CO-B scores, both of which are classified as functionally conserved. DNA elements associated with extended set REPs can exhibit high CO-V scores with or without shared transcription factor binding site (TFBS) motifs.

To quantify CO-V, FUNCODE assumes that samples from different species occupy a similar tissue or cell type space. However, it does not require perfect sample matching or prior information about matched sample pairs. It initially identifies homologous DNA element pairs based on sequence alignment^20^ and then extracts functional genomic data for these pairs. The extracted data are used as features to embed samples from both species into a common low-dimensional space. In the embedding, mutual nearest neighbors within a specified distance between the two species are identified as the *in silico* matched samples^21^, which are then used to score CO-V of each DNA element pair via weighted Spearman’s correlation statistics of the activities across matched samples (**Fig. 1c**). It is worth noting that once the *in silico* matched samples are established, they can be used to score any pair of DNA elements, extending beyond the original set of sequence-aligned element pairs used for creating the matching.

Compared to the conventional approach that analyzes conservation based on manually matched samples between species, FUNCODE’s *in silico* sample matching offers multiple advantages. First, manual sample matching can be error-prone due to incomplete prior knowledge and variations in species-dependent sample collection and experimental procedures. FUNCODE addresses this by learning sample matching directly from the data. Second, manual matching becomes impractical when dealing with a large number of new samples, whereas FUNCODE easily scales to accommodate new samples. Third, FUNCODE’s scalability and ability to align samples without requiring exact matches enable it to include a more extensive set of samples, resulting in more robust and accurate conservation scores than methods that rely on a small number of manually matched samples.

Unlike CO-V, CO-B recognizes element pairs that are consistently active across almost all tissue or cell types in both species, such as promoters and enhancers of housekeeping genes. Computing the correlation may not capture this type of conservation (**Fig. 1b**). To quantify CO-B, we use a statistic inspired by the h-index^22^, which does not require sample matching (**Methods**). FUNCODE annotates DNA element pairs based on their sequence homology, gene homology, CO-V, and CO-B (**Fig. 1d**).

### Quantifying regulatory elements conservation between the human and mouse genomes

We applied FUNCODE to assess the functional conservation of regulatory elements between human and mouse using ENCODE data, focusing on DNase I hypersensitive sites (DHSs) as indicators of regulatory regions^23^. We scored 182 million human-mouse regulatory element pairs (REPs), divided into a core set and an extended set.

The core set, comprising 2.24 million REPs with sequence homology identified via pairwise BlastZ^20^ alignment, included 1.07 million pairs of human and mouse DHSs, 0.87 million pairs of human DHSs aligned with unannotated mouse sequences, and 0.30 million pairs of mouse DHSs aligned with unannotated human sequences (**Methods**). The core set represents 52% of the 3.58 million human DHSs and 61% of the 1.80 million mouse DHSs. For the core set, both CO-V and CO-B scores were computed.

The extended set comprises 180 million REPs identified based on gene homology — pairs of human and mouse DHSs located near orthologous genes but not necessarily aligned by sequence (**Fig. 1a, Methods**). They were considered since enhancer turnover and transcription factor binding site (TFBS) turnover events can result in functionally equivalent regulatory elements with low sequence conservation^11,24,25^. For this set, a median of 12,152 REPs was analyzed for each of the 16,468 human-mouse orthologous gene pairs. We computed CO-V scores for this set since correlated signals may indicate functionally similar elements created by turnover events, irrespective of sequence homology. CO-B was not computed for the extended set because pairs with high baseline activities inherently show high CO-B. Given the larger number of pairs lacking sequence homology, cataloging elements with high baseline activities by species is more efficient and practical. Each REP in the extended set was also annotated based on whether it shared similar transcription factor binding motifs between the human and mouse elements (**Methods**).

We scored functional conservation of each REP for four data modalities, including chromatin accessibility (CA) and H3K4me1, H3K4me3, and H3K27ac histone modifications, using *in silico* matched sample pairs (**Table S1**) identified from a total of 2,595 experiments (**Methods**, **Fig. S1-2, Table S2**). These scores, collectively called FUNCODE scores, are accessible via the ENCODE data portal (**Table S3**). We also developed a web interface to query human or mouse genomic elements by coordinates to retrieve their paired elements and conservation scores (**Fig. S3**).

The FUNCODE scores for REPs allowed us to quantify the conservation of individual human or mouse regulatory elements. Each sequence-aligned element corresponds to one REP in the core set and can be scored using the FUNCODE score of the REP. These were denoted as CO-(V/B)-(CA/H3K4me1/H3K27ac/H3K4me3) scores. Each element in the extended set can be associated with multiple REPs based on gene homology, and it was scored using the maximum FUNCODE score of all associated REPs, denoted as GH-CO-V-(CA/H3K4me1/H3K27ac/H3K4me3). These scores are available as genome browser tracks. For instance, **Fig. 2a-c** shows CO-V-CA, CO-B-CA, and GH-CO-V-CA scores at three loci. The high CO-V-CA in **Fig. 2a** indicates conservation in variable activities of the sequence homology pair. The high CO-B in **Fig. 2b** suggests conserved constitutive activity of the sequence homology pair. In **Fig. 2c**, two pairs of elements with sequence homology are presented; however, neither pair shows functional conservation, as indicated by the CO-V-CA and CO-B-CA scores. Yet, these human and mouse elements also had gene homology, resulting in four element pairs in the extended set. Among these, one pair (the leftmost and rightmost) exhibited high CO-V based on correlated tissue- or cell-type-specific activities, leading to high GH-CO-V scores for the two elements. This high GH-CO-V pair also shared similar TFBS motifs from transcription factors, including SOX4, MEIS2, and MYOG motifs, suggesting it is a potential evolutionary turnover event.

**Fig. 2.**
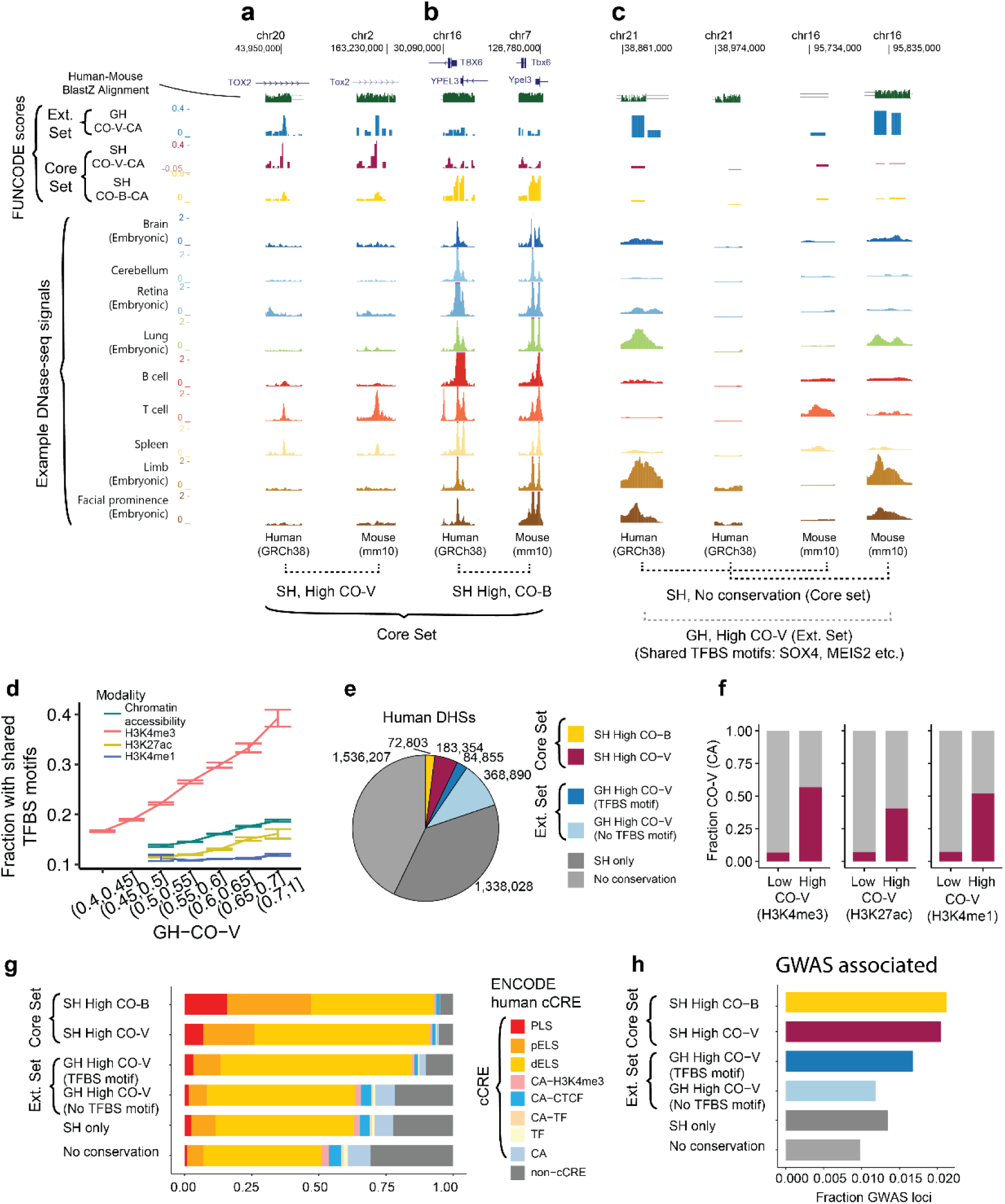
Example FUNCODE score tracks and characteristics of functional conserved elements. (**a**) Genomic tracks displaying a pair of sequence-aligned DNA elements in the intron of the TOX2/Tox2 gene with conserved tissue- or cell-type-specific chromatin accessibility (high CO-V-CA). Highly concordant chromatin accessibility (CA) was observed in T cells and spleen tissue in both species. (**b**) Genomic tracks displaying a pair of sequence-aligned DNA elements near the promoter of TBX6/Tbx6 gene with conserved baseline CA (high CO-B-CA). (**c**) Genomic tracks displaying two pairs of sequence-aligned DNA elements (core set) in intergenic regions with low FUNCODE conservation (low CO-V-CA and low CO-B-CA). CA signals were only observed in embryonic lung, limb, and facial prominence for either human or mouse. Both human elements had gene homology with both mouse elements (four pairs in the extended set), among which a pair of high GH-CO-V-CA elements was identified. The element pair also shared similar TFBS motifs. (SH: sequence homology, GH: gene homology, TFBS: transcription factor binding site) (**d**) Lineplots showing the fraction of REPs that share TFBS binding motifs between human and mouse genomes in bins of GH-CO-V scores. REPs with significant GH-CO-V scores but do not have sequence homology were included. Color represents data modality. Error bars are Mean+/-SEM. REPs with higher GH-CO-V scores were also more likely to share TFBS motifs. (**e**) Pie chart showing the fraction of significant conserved elements among all human DHSs. Color indicates four different conservation categories; Pairs with both high CO-V and high CO-B conservation are categorized as high CO-V. For the amount of overlap, see **Fig. S4**. Similar analysis for mouse is presented in **Fig. S6**. (**f**) Stacked barplot showing the overlap of high CO-V elements defined based on histone modifications with those defined based on CA. (**g**) Stacked barplot showing the enrichment of human candidate Cis-Regulatory Elements (cCRE) among different conservation categories. PLS: promoter, pELS: proximal enhancer, dELS: distal enhancer. TF: transcription factor. (**h**) Barplot showing the fraction of elements with GWAS variants among different conservation categories.

### Characteristics of functionally conserved regulatory elements

By comparing to random expectation, we identified 192,938 and 107,271 REPs with statistically significant conservation in variable activity (high CO-V, any data modality) and baseline activity (high CO-B, any data modality), respectively, in the core set (**Fig. S4, Methods**, FDR ≤10%). In the extended set, we identified 3.07 million REPs with significant conservation in variable activity (GH high CO-V, FDR ≤25%, the more relaxed FDR cutoff was due to lower signal-to-noise ratio), among which 395,390 pairs also shared similar TFBS motifs (**Fig. S5, Methods**). Among all the significant REPs with gene homology but without sequence homology, those with higher GH-CO-V scores were more likely to share TFBS motifs, indicating that they were candidate evolutionary turnover events (**Fig. 2d**).

Across the human genome, the significant REPs in the core set identified 256,157 human DHSs (7.2%) as conserved, and significant REPs in the extended set identified an additional 453,745 DHSs (12.7%) as conserved (**Fig. 2e**). Conserved elements defined by chromatin accessibility and histone modifications exhibited substantial overlap (**Fig. 2f, S6**). For example, 40.5% of high CO-V-H3K27ac were also high CO-V-CA, in contrast to only 7.0% of low CO-V-H3K27ac elements. Comparing conserved and non-conserved DHSs, those with high FUNCODE scores showed greater enrichment of the latest ENCODE candidate cis-regulatory elements (cCREs)^26^ [ENCODE tracking# ENC4P02] (**Fig. 2g**). ENCODE cCREs were curated by integrating multiple data modalities without using functional conservation information. Compared to DHSs exhibiting sequence and functional conservation (core set), DHSs lacking both sequence homology and functional conservation were 5.92 times more likely to be non-cCREs (30.7% vs. 5.2%). DHSs without sequence conservation but with gene homology and functional conservation (extended set) had cCRE proportions similar to or higher than those with sequence conservation but without functional conservation. Within the extended set, DHSs with shared TFBS motif(s) had a higher cCRE proportion than those without (**Fig. 2g**).

We observed distinct cCRE category distributions across different conservation categories. DHSs with sequence homology and high CO-B were more enriched in promoter-like (PLS) and proximal enhancer-like (pELS) elements (47.2% PLS or pELS). In comparison, those with high CO-V were rich in distal enhancer-like elements (dELS) (65.3% dELS) (**Fig. 2g**). Similar results were observed in mouse (**Fig. S6**). Conserved elements also showed higher enrichment in disease-associated GWAS variants compared to non-conserved backgrounds (**Fig. 2h**). Among DHSs with gene homology and high CO-V, those that share TFBS motifs also showed higher enrichment in cCRE and GWAS variants (**Fig. 2g,h**). Enrichment analysis of gene ontology terms using conserved elements showed significant association with essential biological processes such as developmental processes, regulation of transcription, and nucleic acid metabolic processes (**Table S4**).

### Regulatory element conservation is predicative of gene expression conservation

We asked whether high FUNCODE scores of regulatory elements were predictive of conserved expression of their target genes. For this purpose, we computed gene expression conservation scores (CO-V-GE and CO-B-GE) for all human-mouse ortholog gene pairs (**Table S5**) by applying FUNCODE to 742 human and mouse polyA+ and total RNA-seq datasets from ENCODE (**Fig. 3a-d, Table S2**). Regulatory elements were linked to their target genes via the activity by contact (ABC) model predictions in the human genome and by proximity in the mouse genome^27,28^ (**Methods**). When human and mouse elements in a REP were linked to a pair of orthologous genes, the gene pair is identified as the REP’s consistent target (**Fig. 3e**). We found that the CO-V and CO-B scores of a REP were significantly higher if the REP had at least one such consistent target (**Fig. 3f, S7**).

**Fig. 3.**
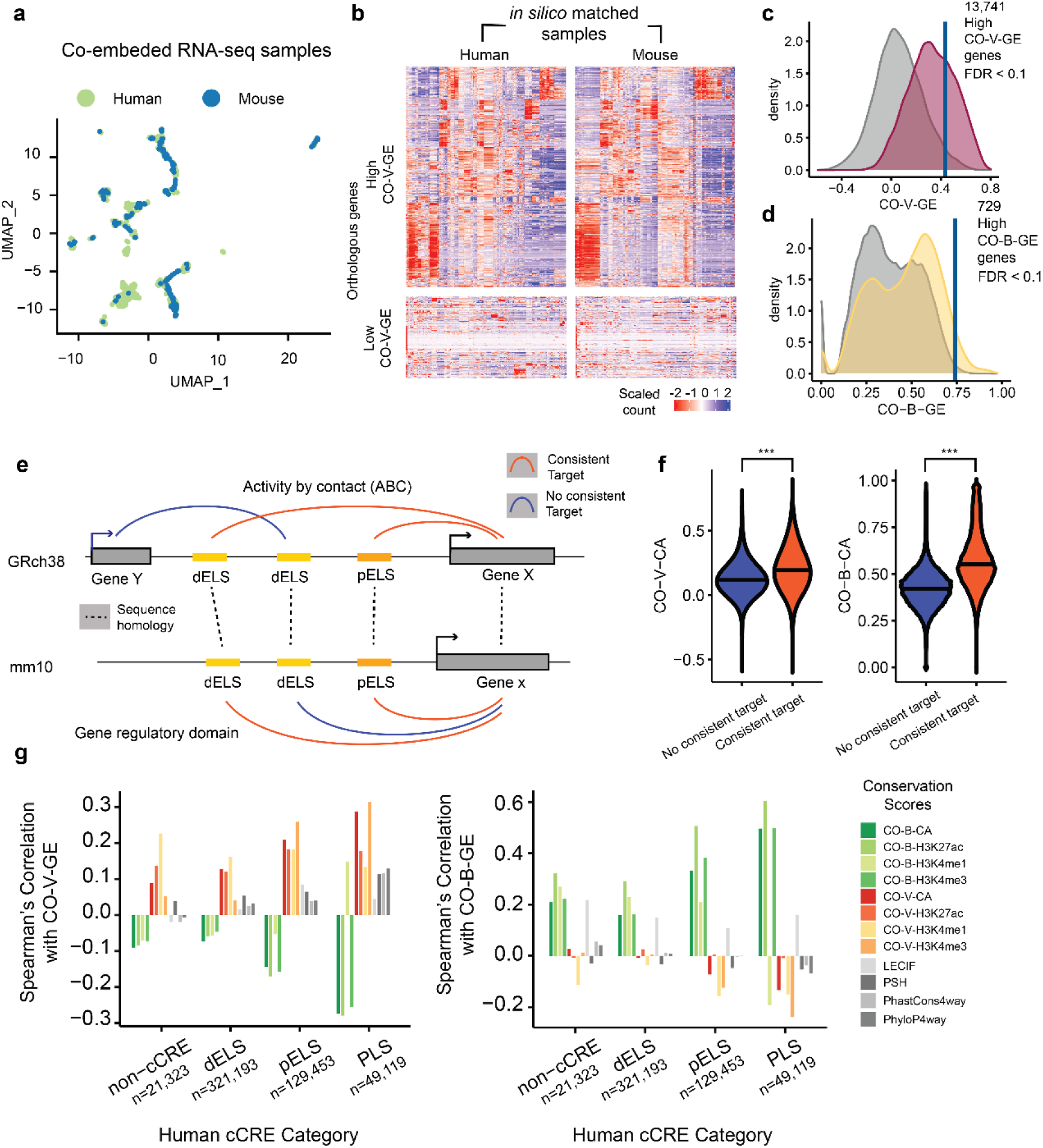
FUNCODE regulatory element conservation scores predict gene expression conservation. (**a**) Scatterplot showing co-embedded human and mouse RNA-seq data. *in silico* matched samples were identified in the common space, facilitating the calculation of gene expression conservation scores (CO-V-GE, CO-B-GE). (**b**) Heatmaps showing example gene expression patterns with either high or low CO-V-GE scores. (**c**) Density plot showing the distribution of CO-V-GE scores of all orthologous genes. The null distribution (created by random pairing of human and mouse orthologous genes) is depicted in gray. A vertical line indicates the significance cutoff at a 10% false discovery rate (FDR). (**d**) Density plot showing the distribution of CO-B-GE scores. The null distribution is colored gray, and a vertical line marks the 10% FDR significance cutoff. (**e**) Schematic showing the categorization of regulatory element pairs based on whether a consistent candidate target gene can be identified. Elements were associated with genes using Activity by Contact model predictions in the human genome and proximity in the mouse genome. Color indicates if the associations were consistent, determined based on whether a pair of orthologous target genes exist. (**f**) Violin plots showing the distributions of CO-V-CA (left) and CO-B-CA (right) scores of the two categories in (**e**). Pairs with a consistent target had higher CO-V-CA and CO-B-CA scores compared to pairs without. ***: Wilcoxon’s rank sum test, *p* < 0.001. (**g**) Barplots showing the Spearman’s correlation (y-axis) between different regulatory element conservation scores and gene expression conservation scores (left: CO-V-GE, right: CO-B-GE) of the consistent target gene pairs across different classes of human cCRE (x-axis). CO-V scores of regulatory elements showed the highest correlation with CO-V-GE and an inverse correlation with CO-B-GE. CO-B scores of regulatory elements were most correlated with CO-B-GE and inversely correlated with CO-V-GE. The strength of these correlations was higher in PLS (promoter), followed by pELS (proximal enhancer), dELS (distal enhancer), and lowest in non-cCRE.

For REPs with at least one consistent target, we computed Spearman’s correlation between the REPs’ FUNCODE scores and the gene expression conservation scores of their closest consistent target genes. We compared FUNCODE scores to a panel of existing genomic conservation scores, including LECIF^17^, phastCons^3^, PhyloP^4^, and percent sequence homology (PSH) (**Methods**). While LECIF was derived from functional genomics data, the other three were sequence-based conservation scores. We focused on regulatory elements in the core set to allow comparison with the sequence-based conservation scores.

Across different cCRE categories, we found that CO-V of regulatory elements was overall positively correlated with CO-V-GE of their target genes and negatively correlated with CO-B-GE of the genes. Similarly, CO-B of regulatory elements was positively correlated with CO-B-GE and negatively correlated with CO-V-GE (**Fig. 3g**). Moreover, FUNCODE scores of regulatory elements more accurately predicted the conservation of functional outcome (gene expression) than other existing scores (**Fig. 3g, S7**). These results suggest that FUNCODE scores of regulatory elements were predictive of gene expression conservation, and the mode of regulatory element conservation (CO-V or CO-B) was also largely consistent with the mode of gene expression conservation.

### FUNCODE better supports cross-species knowledge transfer

FUNCODE can be utilized to determine if findings in one species are likely transferable to another. To benchmark FUNCODE’s performance, we compared it with the panel of existing conservation scores. To allow comparison with sequence-based conservation scores, we again focused on DNA elements that were sequence-alignable between human and mouse genomes (core set).

We first compared different scores’ ability to determine if differential regulatory activities between a pair of new tissue or cell types observed in one species can be transferred to another. We conducted a cross-validation (CV) study in which FUNCODE scores computed using training samples were used to rank differential signals in previously unseen tissue or cell types (**Methods**). Among all conservation scores, FUNCODE CO-V scores most effectively identified transferable differential activities, demonstrating a substantial increase in accuracy (**Fig. 4a, S8a**). Since CO-V required sample matching, we also compared FUNCODE scores derived from the *in silico* matched samples to those derived from manually matched samples (**Table S6**). Notably, CO-V scores computed based on *in silico* matched samples (CO-V) surpassed manually matched samples (CO-V-Manual), underscoring the benefits of the unsupervised approach, where increased power is achieved by generating more matched pairs.

**Fig. 4.**
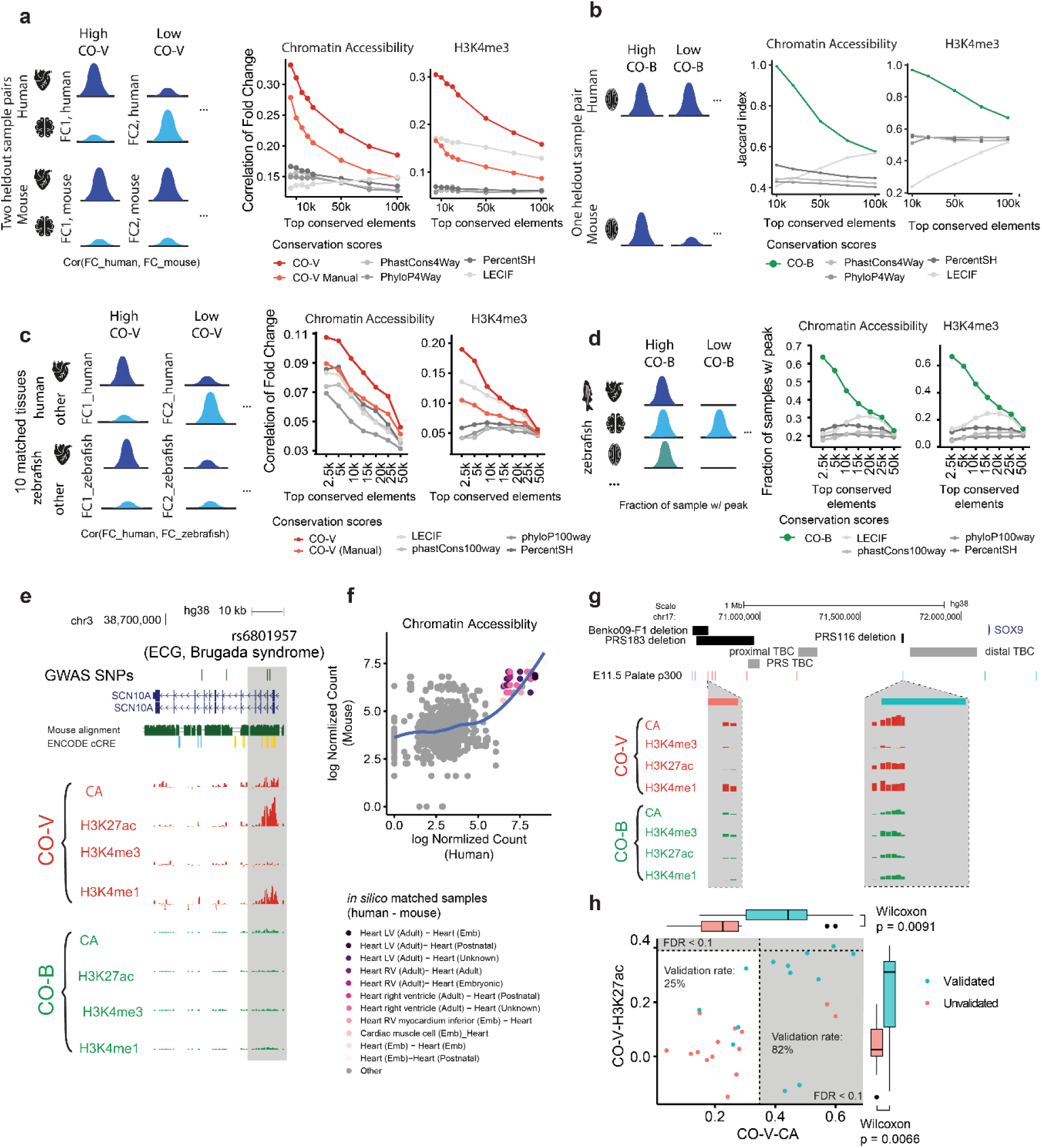
FUNCODE facilitates cross-species knowledge transfer and helps screen human disease-linked regulatory elements. (**a**) Line plots showing conservation of differential activities between new tissue or cell types (as measured by correlation of human and mouse fold changes, y-axis) at different conservation score cutoffs (x-axis). The schematic shows two element pairs with high or low conservation of differential activities. The CO-V score in each panel is the CO-V score specific to the data modality. (**b**) Line plots showing conservation of baseline activities in new tissue or cell types (as measured by the Jaccard index on the y-axis) at different conservation score cutoffs (x-axis). CO-B scores outperformed other conservation scores. The CO-B score in each panel is the CO-B score specific to the data modality. (**c**) Line plots showing conservation of tissue-specific activities between human and zebrafish samples (as measured by correlation of human and zebrafish fold changes, y-axis) at different human-mouse conservation score cutoffs (x-axis). CO-V score outperformed CO-V-Manual and other conservation scores. The CO-V score in each panel is the CO-V specific to the data modality. (**d**) Line plots showing conservation of baseline activities (as measured by the fraction of zebrafish samples with an overlapping peak, y-axis) at different human-mouse conservation score cutoffs (x-axis). CO-B score outperformed other conservation scores. The CO-B score in each panel is the CO-B specific to the data modality. (**e**) Genomic tracks displaying a locus associated with Brugada syndrome. High CO-V-CA, CO-V-H3K27ac and CO-V-H3K4me1 scores were observed. (**f**) Scatterplot showing tissue-specific activities in the heart in both human (x-axis) and mouse (y-axis), which underpins high CO-V scores in the shaded region in (**e**). (**g**) Genomic tracks displaying regions associated with the Pierre Robin sequence of Campomelic dysplasia. Known genetic variations (black and gray) associated with the diseases and p300 ChIP-seq peaks in the E11.5 mouse palate were displayed. Zoomed-in views show FUNCODE scores at two specific p300 peaks. The left peak (red) had low CO-V scores and was not validated by reporter assay in vivo. The right peak (cyan) showed high CO-V scores and was validated. TBC: translocation break clusters. (**h**) Scatterplot showing CO-V-CA (x-axis) and CO-V-H3K27ac (y-axis) scores for elements that overlapped the p300 ChIP-seq peaks in (**g**). Higher CO-V-CA and CO-V-H3K27ac scores were observed for elements that were validated in vivo (cyan vs. red). The grey area indicates regions with significantly high CO-V scores.

Similarly, we compared different methods’ ability to identify conserved baseline activities in new tissue or cell types using another CV analysis (**Methods**). Among all methods, FUNCODE CO-B scores computed from the training set were most effective at identifying high baseline activities transferable across species in the test set, as indicated by the largest Jaccard index characterizing the cross-species transfer consistency (**Fig. 4b, S8b**).

We further reasoned that elements highly conserved between human and mouse would likely remain conserved over extended evolutionary periods. FUNCODE scores defined between these species could thus facilitate knowledge transfer to more distantly related species. To investigate this, we utilized zebrafish ENCODE data^7^, focusing on elements alignable across all three species. Human-mouse FUNCODE scores and other conservation scores were used to rank these elements, and we assessed the performance using ten tissue samples shared between human and zebrafish datasets. Overall, elements highly conserved between human and mouse based on FUNCODE would likely remain conserved in zebrafish. Elements identified with high human-mouse FUNCODE CO-V scores showed a greater Spearman correlation between human and zebrafish for differential signals between two samples. Human elements with high human-mouse FUNCODE CO-B scores also showed higher baseline activities in zebrafish. FUNCODE scores derived from *in silico* matched samples achieved the highest performance, again outperforming manual sample matching (**Fig. 4c, d, S8c, d, Methods**).

Together, these results showed that FUNCODE substantially outperformed other conservation scores for identifying evolutionary conserved regulatory activities and transferring knowledge across species.

### FUNCODE as a resource for prioritizing human disease-associated genetic variants

With its ability to facilitate knowledge transfer, FUNCODE can serve as a tool to guide the development of mouse models for human diseases. We examined previous studies where mouse models were used to validate the function of enhancers identified in human genetics studies. We found that enhancers validated in mouse often had high FUNCODE scores. For example, an enhancer with an SNP associated with Brugada syndrome, a disease with an increased risk of fatal cardiac arrhythmias, was examined (**Fig. 4e**). The enhancer, located in the intron of SCN10A, regulates the nearby cardiac voltage-gated sodium channel gene SCN5A (located 47kb downstream). Previously, a loss-of-function mutation in the SCN5A coding region has been linked to Brugada Syndrome and found in 15-30% of cases^29^. This enhancer was studied in a transgenic mouse model, and its deletion decreased SCN5A gene expression^30^. Indeed, the enhancer had high CO-V scores for CA, H3K27ac, and H3K4me1. Consistent with the conservation, the enhancer displayed high open chromatin activities in embryonic and adult heart samples for both human and mouse samples (**Fig. 4f**).

To systematically demonstrate how FUNCODE can facilitate building mouse models, we analyzed a previous study of the Pierre Robin sequence of Campomelic dysplasia, a disease with distinctive facial features such as a cleft palate, glossoptosis, and micrognathia^31^. Genetic screening in human patients has identified deletion, translocation, and duplications upstream of the SOX9 gene (**Fig. 4g**). Further ChIP-seq experiments targeting p300 proteins in E11.5 mouse craniofacial tissues have identified candidate regulatory elements, which then underwent functional validation by reporter assays. Four of the ten peaks tested showed craniofacial enhancer activity in vivo at E11.5^32^. We compared the CO-V-CA and CO-V-H3K27ac scores for all regulatory elements that overlapped any of the p300 ChIP-seq peaks tested, finding that elements in peaks whose activities could be validated in vivo showed significantly higher CO-V-CA and CO-V-H3K27ac scores. If these regulatory elements were stratified based on FUNCODE scores, the conserved elements would have a much higher validation rate (82%) compared to the non-conserved elements (25%) (**Fig. 4h**), demonstrating that FUNCODE can be used to prioritize candidate elements for building animal models.

### FUNCODE for de novo functional annotation

FUNCODE can facilitate de novo annotation of regulatory elements overlooked in single-species analyses. For example, among the 268,447 conserved REPs (high CO-V or high CO-B) in the core set, 36,144 (13.5%), 47,731 (17.8%), and 20,871 (7.8%) were not currently annotated as mouse cCREs, DHSs, or both (**Fig. 5a, S9a-d**). The functional conservation suggested that many of these unannotated mouse sequences were likely functional regulatory elements. Supporting this hypothesis, a higher proportion (87.8%) of these unannotated mouse elements with high CO-V scores aligned with human cCREs, as compared to the alignment rate of currently annotated mouse elements with low CO-V (76.6%) (**Fig. 5b, S9e, f**). Similar trends were observed in CO-B scores (**Fig. S9g, h**). Moreover, we examined the conservation derived from histone ChIP-seq data targeting H3K4me1, a marker for active and primed enhancers^33^ that is not used by ENCODE for cCREs or DHSs annotation^26^ [ENCODE tracking# ENC4P02]. We found that unannotated elements with high CO-V scores (defined by CA, H3K4me3 or H3K27ac, excluding H3K4me1) exhibited a similar level of CO-V-H3K4me1 as annotated cCREs and DHSs with high CO-V scores, but showed a higher level of H3K4me1 conservation compared to annotated cCREs and DHSs with low CO-V scores (**Fig. 5c, S9i, j**). Similarly, unannotated elements with high CO-B scores (defined by CA, H3K4me3 or H3K27ac, excluding H3K4me1) also exhibited high baseline conservation in H3K4me1 (**Fig. S9k, l**). These results suggest that the conserved unannotated elements are likely real regulatory elements. Transferring annotations of FUNCODE-conserved elements across species yielded 36,144 putative new cCREs and 47,531 new DHSs in the mouse genome. Similarly, we identified 6,561 putative new cCREs and 12,957 DHSs in the human genome. The smaller numbers for human were due to more extensive data collection compared to in mouse. These FUNCODE-annotated new cCREs or DHSs were made available as an open resource (**Table S7**).

**Fig. 5.**
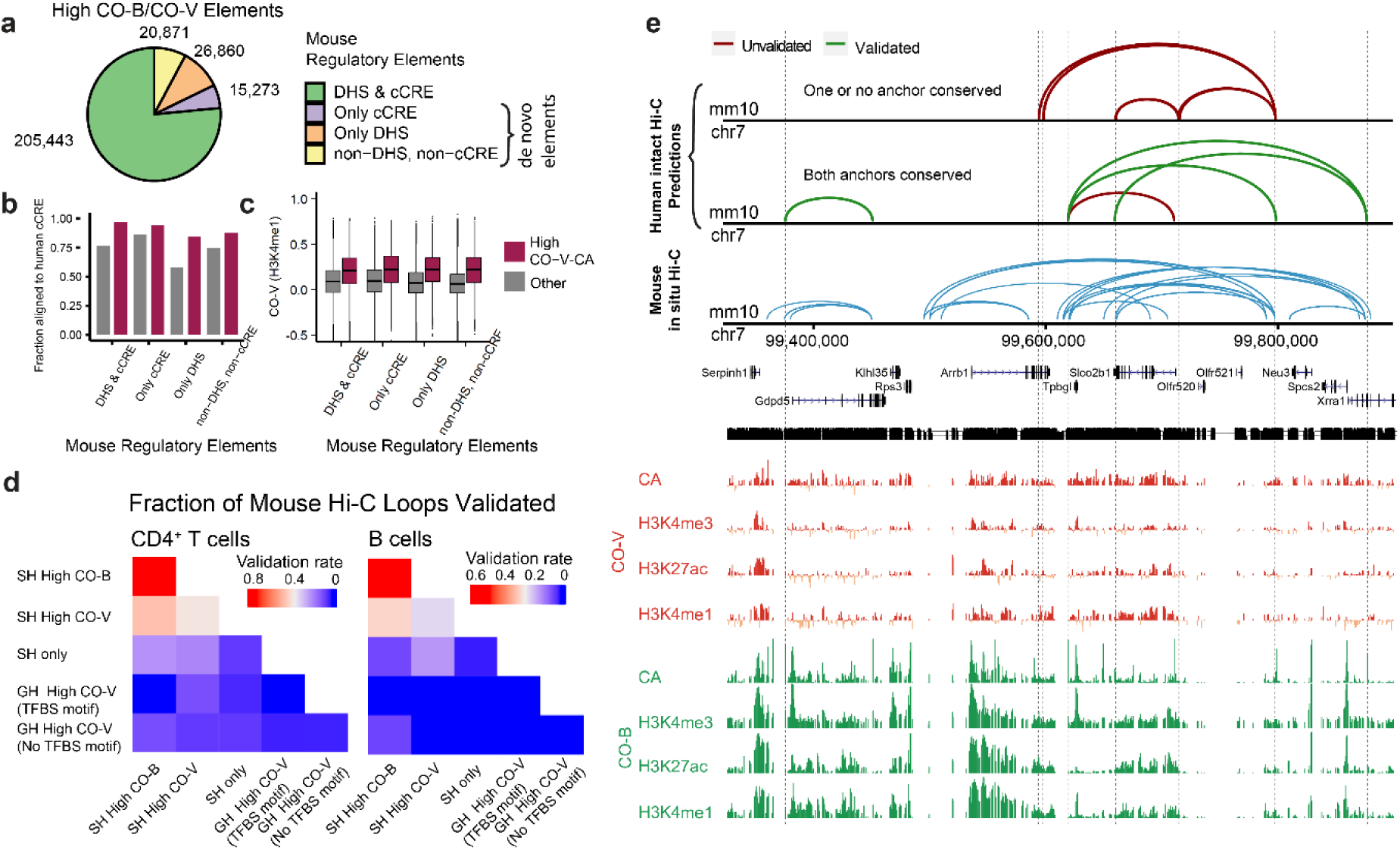
FUNCODE enables de novo functional annotations and transfer of chromatin interactions across species. (**a**) Pie chart breaking down the current annotations of the conserved elements called by FUNCODE in the mouse genome. (**b**) Barplot showing alignment rate of mouse DNA elements to human cCRE (y-axis) for different categories in (**a**) (x-axis). Color indicates high CO-V elements versus others. Conserved (high CO-V) but unannotated mouse DNA elements had similar rates to conserved and annotated elements, and both showed higher rates compared to annotated but non-conserved elements. (**c**) Boxplot showing the distribution of CO-V-H3K4me1 scores (y-axis) for different categories in (**a**) (x-axis). Conserved (high CO-V, defined without using H3K4me1 ChIP-seq data) but unannotated mouse elements had similar CO-V-H3K4me1 scores compared to conserved and annotated elements, and both had higher scores compared to annotated but non-conserved elements. (**d**) Heatmaps showing the validation rates of human chromatin interactions transferred from mouse Hi-C data in CD4^+^ T cells (left) or B cells (right). Each cell represents a unique combination of REPs for mapping the loop anchors across species. For example, in CD4^+^ T cells, the highest validation rates by human Hi-C data were observed when both loop anchors were aligned by sequence homology (SH) and high CO-B conserved (first row, first column). (**e**) Genomic tracks showing mouse contact loops from in situ Hi-C data (bottom) or predicted mouse contact loops transferred from human ENCODE Hi-C (top and mid), all for CD4^+^ T cells. The top track has one or no loop anchors conserved (High CO-V/CO-B). The middle track has both loop anchors conserved. Green contact loops were validated by the mouse in situ Hi-C data, and red loops were unvalidated. CO-V and CO-B score tracks are shown below. Predictions based on conserved loop anchors have a higher validation rate.

We also explored the potential of FUNCODE in identifying 3D chromatin interactions through cross-species knowledge transfer, leveraging ENCODE’s Hi-C data resource. The latest ENCODE Hi-C data identified loops associated with individual regulatory elements^34^ [ENCODE tracking # ENC4P85]. By mapping both loop anchors across species via REPs, we transferred Hi-C loops between human and mouse genome and focused on transferred loops with a CTCF binding motif on at least one of the loop anchors. To benchmark the transfer accuracy, we assessed human and mouse Hi-C datasets of CD4^+^ T cells and B cells. We evaluated transfer accuracy from mouse to human by computing the fraction of loops transferred from mouse that can be validated using human Hi-C data in the corresponding cell type (**Table S2, Methods**). Because the mouse data was generated using an older and less sensitive protocol^35^ [ENCODE tracking # ENC4P10], performing evaluation in the reverse direction (human-to-mouse) would underestimate the validation rate. Loop anchors exhibiting both sequence and functional conservation (SH high CO-B or CO-V) exhibited higher validation rates than those with only sequence conservation (SH only). The highest validation rates were achieved for constitutively active loop anchors exhibiting both sequence and functional conservation (SH high CO-B). For instance, when mapping mouse CD4^+^ T cell loops to human, the validation rate was 76.7%. As a negative control, we also examined loop anchors that are near homologous genes but did not align by sequence (gene homology REPs). As expected, the validation rate was much lower (3.4%, **Fig. 5d, S9m, n**).

Because ENCODE generated far more Hi-C data in human than in mouse, transferring ENCODE human Hi-C data to mouse could provide a valuable resource for linking regulatory elements to genes in mouse. As a proof-of-principle, we transferred data from eight human primary immune cell types to the mouse genome. To maximize the accuracy of the transfer procedure, we only retained loops where both anchors exhibited sequence and functional conservation. In total, this procedure identified 37,968 high-confidence transferred Hi-C loops, and we labeled them as FUNCODE-predicted mouse contact loops (**Fig. 5e, Table S8**). We estimated the false discovery rate for these predictions using the control procedure described in the preceding paragraph, examining loops anchors near homologous genes but without sequence alignment. This resulted in estimated FDR ranging from 2.3% in B cells and 4.9% in CD4^+^ T cells (**Methods**). This application of FUNCODE to chromatin interaction data demonstrates the utility of functional conservation scores in transferring complex data modalities across species, bridging gaps in available experimental data.

### FUNCODE enhances integrative analysis of single-cell omic data across species

Integrative analysis of human and mouse single-cell omics data typically involves selecting features with sequence homology across species and highly variable activities across single cells. These features are then used to embed cells from different species into a common space. FUNCODE can improve this integration by further focusing on features with conserved tissue- or cell-type-specific activities (high CO-V) across species (**Methods**). To demonstrate, we integrated published human and mouse sciATAC-seq data from four tissues (lung, liver, heart, large intestine)^36,37^ using standard protocols, but with an added step of selecting features (sequence-aligned regulatory elements) based on their CO-V scores. We focused on CO-V scores since integration aims at mapping similar cell types across species, and high CO-B or non-conserved elements are not informative for this purpose.

To assess the integration efficacy, we transferred known cell type labels from human to mouse, and evaluated the accuracy of the label transfer using the known mouse cell type labels, as measured by the Adjusted Rand Index (ARI). Remarkably, using an equal number of features for integration, those filtered by FUNCODE’s CO-V-CA scores achieved substantially higher ARI compared to those filtered by other existing scores, including a control method where no conservation filtering was applied (**Methods, Fig. 6a-d**, **S10**). For instance, when features were filtered by CO-V-CA score, most mouse hepatocytes identified by label transfer were indeed hepatocytes. This success rate dropped markedly with other methods. A similar trend was observed for mouse endothelial cells (**Fig. 6e**).

**Fig. 6.**
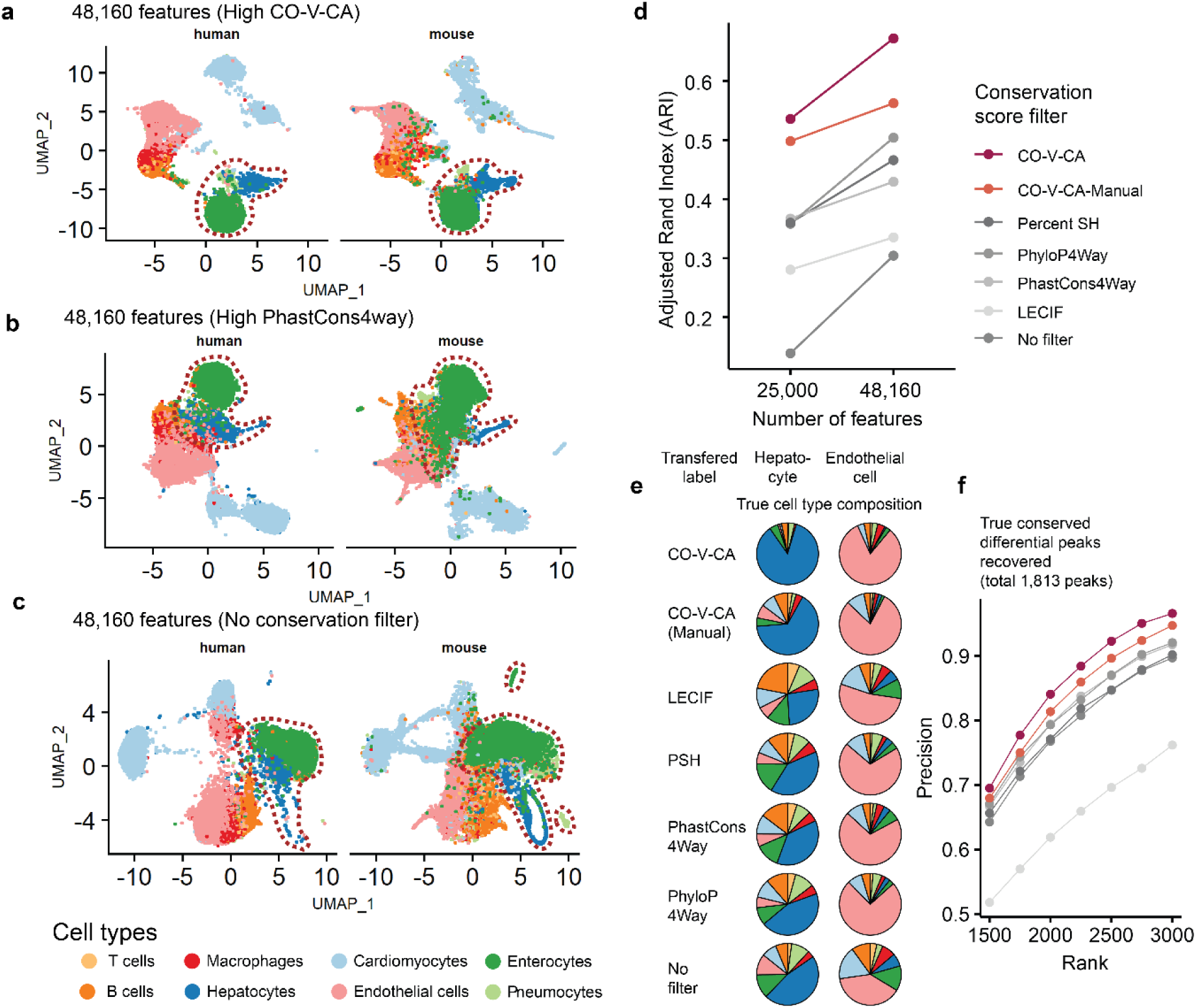
FUNCODE scores facilitate single-cell genomics data integration across species. (**a**) Scatterplot showing UMAP embedded single cells from human and mouse sciATAC-seq datasets after cross-species integration. Human and mouse cells were shown in separate panels. Color indicates cell types labeled in the original publications. 48,160 features (sequence-aligned element pairs) with the highest CO-V-CA scores were used for the integration. Red-brown dashed outlines highlight hepatocytes, enterocytes, and pneumocytes, demonstrating improved cross-species mapping and clearer separation from other cell types when the high CO-V-CA feature filter was applied. (**b**) Same as (**a**), but the features with the highest PhastCons4Way scores. (**c**) Same as (**a**), but no filtering of features with conservation scores was applied. (**d**) Lineplot showing label transfer accuracy (as measured by adjusted rand index, y-axis) for integration at different conservation score cutoffs (x-axis). Color indicates different conservation score filtering during integration. The standard integration approach currently in use does not apply a conservation filter (No filter). (**e**) Pie charts showing the compositions of transferred human cell type labels for mouse hepatocytes and endothelial cells. Rows indicate different conservation score filtering during integration. Color indicates cell types. (**f**) Lineplot showing the fraction of true conserved differential element pairs between hepatocytes and endothelial cells recovered among the inferred pairs. The inferred pairs were defined under the assumption that mouse cell types were unknown and had to be transferred from the human dataset. Color indicates different conservation score filtering during integration.

This improved integration enables more accurate comparative analyses of regulatory mechanisms between species. For instance, consider identifying differential chromatin accessibility between hepatocytes and endothelial cells conserved between human and mouse samples. First, we established a set of ground truth conserved regulatory element pairs using differential signals computed from known human and mouse cell type labels. We then sought to determine how effectively these conserved differential regulatory signals could be identified without prior knowledge of mouse cell types, relying instead on data integration. When filtering with different conservation scores, human cell type labels were transferred to mouse cells based on their respective integration results, and differential chromatin accessibility was computed between the two cell types in each species in each case. The regulatory elements were then ranked based on their combined fold change in human and mouse, and the recovery rate of ground truth conserved regulatory element pairs was examined (**Methods**). Our findings indicated that the integration based on FUNCODE’s CO-V-CA scores most effectively identified true conserved differential elements. It outperformed integration without using conservation filters (current state-of-the-art) as well as integration based on other existing scores (**Fig. 6f**). Therefore, FUNCODE enables more accurate identification of conserved regulatory signals in cross-species single-cell analysis, potentially overlooked in standard single-cell integration protocols.

## Discussion

In summary, FUNCODE is a framework for scoring cross-species functional conservation of DNA elements based on unsupervised sample matching. This strategy ensures that the matching is objective and negates the need for manual annotation. Data-driven sample matching also enabled more efficient use of existing datasets. Our approach, combined with systematically generated and uniformly processed human and mouse data by the ENCODE consortium effort, produced conservation scores useful in many different tasks. Similar to traditional sequence conservation scores, we showed that FUNCODE scores provide orthogonal information to help annotate functional parts of the genome. Unlike traditional scores, FUNCODE scores have the additional utilities of predicting functional signals of chromatin accessibility, histone modifications, and chromatin interaction in unseen tissue, cell types, or even distantly related species. These abilities are essential for applying conservation scores to help screen human disease-associated elements as candidates for animal models.

Our framework generated a genome-wide catalog of functionally conserved regulatory elements between humans and mice, including potential evolutionary turnover events. This catalog provides a foundational resource for elucidating genome functions, enhancing cross-species analysis, and advancing disease modeling. The FUNCODE catalog of putative turnover events is broad and could be further refined with additional functional data. For example, more advanced sequence models^38,39^ [ENCODE tracking #ENC4P28, #ENC4P40] could be used to further screen for elements that have failed to align due to non-functional mutations.

The flexibility and scalability of FUNCODE means that it can be reapplied when additional samples become available. Existing scores are expected to improve during the process. One potential limitation of unsupervised sample matching is requiring a diverse panel of samples in each species to cover a common space of tissue or cell types. This limitation, however, can be mitigated by the continued growth of functional genomics data. For one, when more data becomes available in other data modalities, new FUNCODE scores can be defined. For the other, while the current work is concentrated on the mouse, the most widely used model organism for which the most complete data was generated, it may also be applied to other species when more data become available, much like the sequence conservation framework can be extended to include more species.

## Methods

### The core set of regulatory elements pairs aligned by sequence homology (SH)

To identify core regulatory element pairs, we obtained the list of DNase I hypersensitive sites (DHSs) for the human GRCh38 genome from the ENCODE data portal (accession: ENCFF503GCK). To map these regulatory regions to the mouse genome, we used pairwise sequence alignment produced by BLASTZ, downloaded from the UCSC genome browser ( https://hgdownload.soe.ucsc.edu/goldenPath/hg38/vsMm10/hg38.mm10.net.axt.gz). Initially, we mapped each summit of the human DHS to the mouse genome. Subsequently, we created a 200 bp window centered around each human summit and its corresponding aligned mouse position to ensure consistency in mapped region widths. This collection of mappings was designated as the ‘forward mapped set’.

Similarly, DHSs for the mouse mm10 genome were retrieved (ENCODE accession: ENCFF910SRW), and the mouse DHSs were aligned to the human hg38 genome. Since mouse DHSs did not have summit points defined, we centered the mapping on the midpoint of the DHSs. The reverse BLASTZ alignment from mm10 to hg38 (https://hgdownload.soe.ucsc.edu/goldenPath/mm10/vsHg38/axtNet) was used to map the midpoint, and 200bp windows were created. The resulting pairings formed the ‘reversed mapped set’.

We then removed duplicates — pairs present in both the forward and reversed mapped sets — by comparing pairs across the sets. A reversed mapped pair was considered a duplicate if both its human and mouse regulatory elements overlapped by more than 50 base pairs with any pair in the forward mapped set. After deleting these duplicates, we combined the remaining reversed mapped pairs with the forward set to finalize the core set of sequence-aligned regulatory element pairs.

### The extended set of regulatory element pairs aligned by gene homology (GH)

To further expand our analysis to pairs of regulatory elements that cannot be sequence-aligned, we identified regulatory element pairs based on gene homology. Using the Mouse Genome Informatics resource from the Jackson Laboratory website (www.informatics.jax.org/downloads/reports/HOM_MouseHumanSequence.rpt), we selected 16,468 orthologous gene pairs. We defined gene regulatory domains for human and mouse genes using Gencode gene annotations from the ENCODE data portal (Human Accession: ENCFF159KBI, Mouse Accession: ENCSR884DHJ). For each gene, the basal domain spanned from 5kb upstream to 1kb downstream of the transcription start site (TSS). These domains were then extended to the nearest gene boundary or up to a maximum of 1MB—the regulatory region associated with a gene included, at minimum, its basal domain.

Regulatory elements within these domains were then paired, with each human element matched to every corresponding element in the homologous mouse gene’s domain, provided a gene homolog existed. This method yielded a median of 12,152 pairs per homologous gene and a total of 1.8×10^8^ candidate pairs for the extended set.

### The panel of existing conservation scores

We compiled a panel of existing conservation scores for comparison with FUNCODE scores; they were processed as follows.

#### PhastCons and PhyloP

PhastCons and PhyloP scores for the human hg38 genome were obtained in bigwig format from the UCSC genome browser (https://hgdownload.cse.ucsc.edu/goldenPath/hg38/). The ‘bigWigAverageOverBed’ tool from UCSC (http://hgdownload.cse.ucsc.edu/admin/exe/) was used to calculate the average scores across each element. The four-way PhastCons and PhyloP scores based on alignments of Homo sapiens, Mus musculus, Galeopterus variegatus, and Tupaia chinensis represent the smallest datasets containing both human and mouse and were shown as representative examples (**Fig. 4**). Those computed from more species yielded consistent results (**Fig. S8**).

#### Percent Sequence Homology (PSH)

For the Percentage of Sequence Homology (PSH), we focused on the part of each pair that can be pairwise aligned and computed the ratio of identical bases in the aligned regions relative to the total number of bases in the human region.

#### LECIF

LECIF score in bigwig format was downloaded for hg19 and mm10 from the GitHub repo (https://github.com/ernstlab/LECIF)^17^. For regions defined on hg38, we converted the coordinates to hg19 using ‘liftOver’. For each region, the LECIF score was averaged.

### Chromatin accessibility and histone modification data processing

We retrieved all human and mouse DNase-seq, ATAC-seq, and Histone ChIP-seq data (targeting H3K4me1, H3K4me3, H3K27ac) from the ENCODE data portal. Detailed experiment listings can be found in **Table S2**. We downloaded read fragments in ‘.bam’ format and quantified the overlap of read centroids with each regulatory element. For paired-end reads, centroids were determined by averaging the 5’ ends of forward and reverse reads, whereas for single-end reads, centroids were the midpoints of reads. We calculated the library size for each sample based on the total number of aligned reads (single-end) or read pairs (paired-end). The library size for each sample was computed as the total number of reads (single-end) or read pairs (paired-end) aligned to the genome. The counts and library sizes of the technical replicates were first averaged for each ENCODE experiment. Then, the counts of each experiment were normalized by dividing by the library size and subsequently multiplying by a factor of 100 million (1×10^8^).

### Standardized variance calculation for regulatory elements

To determine highly variable regulatory elements based on the functional genomics datasets, we calculated standardized variance for each regulatory element. Following a similar routine as the Seurat package (version 3.0)^40^, we started by computing the variance and mean of each regulatory element. Subsequently, a lowess smoothing was applied to the relationship between log-transformed variance and mean. For each regulatory element, the lowess prediction, conditional on its mean expression, is used to derive the expected standard deviation. Each feature was then standardized by subtracting the mean and dividing it by the expected standard deviation. Standardized values exceeding the square root of the total sample count were clipped off to mitigate the influence of outliers. The final standardized variance is the variance of these standardized scores. This procedure is known as the variance stabilizing transformation (vst). Notably, unlike Seurat’s application of VST directly to raw count data, our approach applies VST to data normalized by library size.

### Assay-specific effect correction between DNase-seq and ATAC-seq data

Before conducting the cross-species analysis, adjusting for assay-specific disparities between DNase-seq and ATAC-seq datasets is crucial. Our approach corrects the data in the original high-dimensional space of regulatory elements. It removes assay effects from individual regulatory elements instead of harmonizing in the low-dimensional embedding. To leverage the more abundant DNase-seq experiments, we treated DNase-seq samples as the reference and ATAC-seq samples as the query set. Employing Seurat’s method (version 3.0)^40^, the assay effects were corrected in two steps. The first step identified anchors following canonical correlation analysis (CCA), and the second step integrated the data by computing a correction vector for each sample in the query dataset. 10,000 shared variable features were used during the anchor finding step, and the top 30 canonical variables and five nearest neighbors (k.neighbor) were considered. Ten nearest neighbors were used when filtering anchors (k.filter). Subsequently, the ‘IntegrateData’ function was called to calculate the correction vectors for the ATAC-seq samples, which were the Gaussian kernel weighted differences of the feature vectors between the anchor pairs. The weighting used the top 20 canonical variables and included up to 20 neighbors (k.weight) (**Fig. S1**).

### Unsupervised integration of human and mouse chromatin accessibility data

To establish *in silico* correspondences between human and mouse samples, we adopted the mutual nearest neighbor (MNN) approach initially proposed by Haghverdi et al. The approach has demonstrated efficacy in scenarios where cell state compositions vary between the datasets to be integrated^21^. In our methods, both human and mouse data were initially projected onto a shared low-dimensional space using Canonical Correlation Analysis (CCA). Within this space, we identified *in silico* matched human and mouse sample pairs using the MNN criteria. A pair of samples, one from each species, is considered a mutual nearest neighbor if each is the nearest neighbor of the other in their respective species. This mutual selection helped to ensure that the pairs of samples were genuinely similar in their biological state instead of due to spurious correlation. Assessing cross-species conservation and computing FUNCODE scores will only depend on these *in silico* matched samples afterward. However, the human and mouse samples were further integrated based on these identified anchor pairs for visualization. The integration corrected the data so that the MNN pairs were close to one another after the correction.

The procedure was executed as follows: We began by using the core set of sequence-aligned regulatory elements as the input and excluded element pairs with normalized counts below 20 in either all human or all mouse samples. We then calculated standardized variances for each element in both species separately, as detailed in the previous section. We did an initial filtering step for elements exhibiting high standardized variance in both species. In detail, each element was ranked based on its standardized variance in both species and the human and mouse ranks were added. The top 30%, based on the overall rank, were selected as our initial feature set for integration. We employed the main pipelines in Seurat v3 for dimension reduction and anchor (MNN) identification. The anchor finding procedure is Seurat’s implementation of the mutual nearest neighbor (MNN) procedure. The mutual nearest neighbor pairs were called anchor pairs and were further filtered and scored based on their quality. Seurat was originally designed for single-cell omics data; we tailored the parameters to the sizes of our bulk compendium. Among the initial set of features, the Seurat pipeline further identified the 10,000 top shared variable features for Canonical Correlation Analysis (CCA), and pairwise distances were computed using the top 30 canonical variables. Five nearest neighbors (k.anchor) were used to determine mutual nearest neighbors. Ten nearest neighbors were used when filtering anchors (k.filter), and 30 nearest neighbors were used to score anchors (k.score). The identified anchor pairs are listed in **Table S1**. Finally, we integrated the datasets for visualization and ran Uniform Manifold Approximation and Projection (UMAP)^41^ on the integrated data, as displayed in **Fig. S2a**.

### Unsupervised bridge integration of human and mouse histone ChIP-seq data

To identify *in silico* matched samples for histone modification data, we first transferred Histone ChIP-seq data to the chromatin accessibility data space and subsequently identified *in silico* matched samples across species based on the transferred data. This approach, termed “bridge integration”, capitalizes on the rich chromatin accessibility data available for both species, using it as a pivotal link between them. Bridge integration was applied separately to Histone ChIP-seq data targeting H3K4me3, H3K27ac, and H3K4me1.

To conduct the bridge integration, we first transferred histone ChIP-seq data onto the CA (DNase-seq and ATAC-seq) data space for human and mouse samples separately. We used the data transfer pipeline by the Seurat package^40^ for this step. First, we narrowed down to a set of features that were highly variable for both histone modification and CA. After computing the standardized variance of the Histone ChIP-seq data for all regulatory elements, standardized variance ranks of individual elements were computed and added to the standardized variance ranks based on CA. The top 15% highest ranked regulatory elements were chosen as the cross-modality integration features between histone modification and CA. Next, after Canonical Correlation Analysis (CCA), transfer anchors were identified (Seurat function FindTransferAnchors) based on 30 top canonical variables and eight nearest neighbors (k.anchor), filtered based on 20 nearest neighbors (k.filter) and scored based on 40 nearest neighbors (k.score). Based on these anchors, Histone ChIP-seq data can be transferred onto the CA space (Seurat function TransferData). For each histone modification sample, this was done by finding a set of nearest anchors in the histone ChIP-seq space, and then taking a weighted average of the nearest anchors in the CA space. The weight was determined based on the distances of the sample to the anchors. Five nearest anchors were considered (k.weight) for weighting. The weighted average can be taken for any set of features of the CA samples. Because our goal was to identify pairing across species, we averaged the 10,000 CA features when integrating CA data. Finally, the transferred histone modification samples were concatenated to the original CA samples, and *in silico* sample pairs were identified using the same procedure as the last section.

### Computation of FUNCODE conservation scores

We calculated Conservation (CO) scores using *in silico* matched samples to assess the conservation of DNA elements across human and mouse genomes. For each pair of DNA elements indexed by *i*, we defined *X_i_*= (*x_i_*_1_, *x_i_*_2_, … , *x_iJ_*) as the vector of log normalized counts for human anchors and *Y_i_* = (*y_i_*_1,_*y_i_*_2_, … , *y_iJ_*) as the vector of log normalized counts for mouse anchors. We assume there are *J* total anchors. Additionally, *w* = (*w*_1_, *w*_2_, … , *w_J_*) represents the vector of anchor scores computed during the anchor finding step. The anchor scores were used as weights for incorporating the anchor quality information.

To quantify variable activities, we defined the CO-V as the weighted Spearman’s correlation between *X*_*i*_ and *Y*_*i*_ :

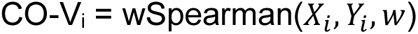

Here, we briefly describe the method for computing the weighted Spearman’s correlation. First, the weighted rank *r_*i*j_* for each element *x_ij_* of the vectors *X_*i*_* was computed where *r_*i*j_* = *a_*i*j_* + *b_*i*j_*. Here *a_*i*j_* is the rank without considering ties and *b_*i*j_* deals with ties. Specifically,

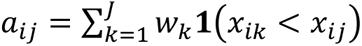

where 1(·) denotes the indicator function. And

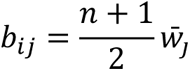

where there are *n* values tied with *x_*i*j_* whose average weights were 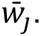 Then, the same procedure is applied to calculate the weighted ranks *s_*i*j_* for each element of *y_*i*j_* in *Y_*i*_*. Finally, the weighted Spearman’s correlation is calculated as the Pearson’s correlation of the weighted rank vectors (*r_*i*_*_1_, *r_*i*_*_2_, … , *r_*i*j_*) and (*s_*i*_*_1_, *s_*i*_*_2_, … , *s_*i*j_*)^42^.

To quantify constitutive activities, we developed the CO-B score, which evaluates both the magnitude and consistency of regulatory element signals across species. A high CO-B score indicates elements with consistently strong activity in both human and mouse, suggesting robust regulatory function across various cell and tissue types. The CO-B score is conceptually similar to the h-index, a metric commonly used in scientific publishing^22^.

We began by calculating the global percentiles of log-normalized signal values across all regulatory elements and samples in both species. By pooling these signal values, we defined *c_k_* and *d_k_* as the *k*-th percentile of these pooled values for human and mouse, respectively. For each *k*-th percentile, we computed the fraction of anchors with signal values exceeding this percentile in human and mouse, denoted as *p*_*ik*_ and *q*_*ik*_:

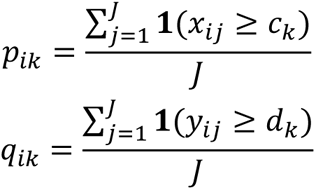

where **1**(⋅) is the indicator function, and *x_*i*j_* and *y_ij_* represent the signal values for human and mouse samples, respectively.

The CO-B score for the regulatory element is defined as the maximum *k* such that *k* percent of the matched samples have signal values exceeding the global *k*-th percentile in both species.

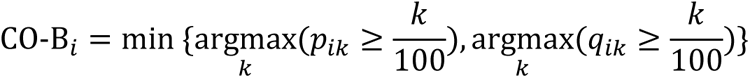

This definition ensures that the CO-B score reflects both signal magnitude and consistency. To illustrate this, consider two examples. In terms of magnitude, an element with all samples at the global 50^th^ percentile will score 0.5, while one consistently at the 30^th^ percentile will score 0.3. This difference reflects the higher magnitude of activity in the first element. Regarding consistency, an element with 80% of samples at the 50^th^ percentile scores 0.5, while another with 40% of samples above the 80^th^ percentile (and the rest below the 40^th^ percentile) scores 0.4. Despite some very high signals, the second element’s lower score reflects its lack of consistency across all samples. Importantly, a high CO-B score can only be achieved when consistently strong signals are observed in both species, underscoring the element’s functional conservation across human and mouse.

### Calling statistically significant pairs of regulatory elements

To rigorously assess the significance of FUNCODE scores, we constructed gene context-dependent null distributions. This involved categorizing human and mouse regulatory elements based on their proximity to the nearest transcription start site (TSS). Specifically, regulatory elements within 200 bp of a TSS were designated as promoter elements, those between 200 bp and 2,000 bp as proximal enhancer elements, and those beyond 2,000 bp as distal enhancer elements. Consequently, each human-mouse regulatory element pair was classified into one of nine possible combinations: promoter-promoter, promoter-proximal, distal-distal, and so on.

To generate null distributions, we randomly paired human and mouse regulatory elements for each category. For instance, to create a null distribution for promoter-proximal pairs, we randomly selected a human promoter element and paired it with a randomly chosen proximal mouse element. This random pairing process was replicated 100,000 times for each of the nine categories.

Subsequently, we calculated CO-B and CO-V scores for all these randomly paired null sets. The empirical p-values were then determined by comparing the CO scores of actual regulatory element pairs against the empirical null distribution of their corresponding category. The p-values of all pairs were further adjusted for multiple testing with the Benjamini Hochberg procedure^43^. A false discovery rate (FDR) cutoff of 0.1 was applied for calling the conserved elements in the core set, and an FDR cutoff of 0.25 was applied for calling significant pairs in the extended set.

### ENCODE candidate Cis-Regulatory Elements (cCRE) version 4 annotations

ENCODE cCRE v4 annotations were collected from the ENCODE data portal^26^ [ENCODE tracking# ENC4P02]. These annotations categorized cCREs into promoter-like sequences (PLS), proximal enhancer-like sequences (pELS), distal enhancer-like sequences (dELS), and elements with CA and H3K4me3 (CA-H3K4me3), elements with CA and CTCF binding (CA-CTCF) and elements with CA and transcription factor binding (CA-TF), elements with only TF binding (TF) and elements with only CA (CA). Elements in the core and extended set were overlapped with each cCRE category and labeled accordingly. In cases where an element overlapped with more than one category, we used the following order to determine the label: PLS > pELS > dELS > CA-H3K4me3 > CA-CTCF > CA-TF > TF > CA. For example, if an element overlapped both a PLS and a pELS, it was labeled as PLS.

### Mapping transcription factor binding site (TFBS) motifs to regulatory elements

The non-redundant CORE motifs were downloaded from JASPAR^44^, which contains 630 human and 106 mouse motifs. Next, all 736 motifs were mapped to the human reference genome hg38 and the mouse reference genome mm10 using the motifmap_matrixscan_genome function in CisGenome^45^. This resulted in a binary motif occurrence matrix (number of elements × number of motifs) for human and mouse, in which a value of one indicates the presence of a motif, and zero value indicates the absence of a motif. Due to the similarity among many transcription factor binding site (TFBS) motifs and their occurrence patterns, we employed non-negative matrix factorization (NMF)^46^ with a low dimensional rank of 25 to reduce 733 motifs to 25 distinct meta-motifs (**Fig. S5e**). NMF was chosen for its ability to maintain the interpretability of latent dimensions. Specifically, let *V*^*T*^ be the motif indicator matrix (number of elements x number of motifs). NMF will identify a coefficient matrix *H*^*T*^(number of elements × number of meta-motifs) and a weight matrix *W*^*T*^(number of meta-motifs × number of motifs) such that *V*^*T*^ = *H*^*T*^ × *W*^*T*^, with the matrix transpose introduced to maintain consistency with the original literature.

Consequently, the motif occurrence matrix *V*^*T*^ (733 columns) is reduced to the meta-motif coefficient matrix *H*^*T*^ (25 columns), with rows of *W*^*T*^ specifying the motif weights for each meta-motif. We further binarized the coefficient matrix *H*^*T*^ by standardizing the meta-motif coefficient vector of each element, considering a DNA element as having a particular meta-motif if its standardized coefficient (z-score) for the meta-motif is above 2.0 (**Fig. S5f**). A pair of human and mouse DNA elements were considered to share TFBS motifs if they have at least one common meta-motif. Identifying shared TFBS motifs allowed for identifying a more confident set of conserved DNA element pairs in the extended set representing candidate evolutionary turnover events.

### NHGRI-EBI GWAS catalog data processing and annotation

All GWAS associations (v1.0) were downloaded from the NHGRI-EBI GWAS catalog^47^. We employed a pruning process based on a previously established method^48^ to refine this data. First, the GWAS SNPs were grouped according to study (PubMed ID) and disease/trait. For each study and disease/trait combination, all SNPs were ranked by their p values. Next, each SNP was added to the final list by checking if there was an SNP within 5,000 bp in the current list; that is, an SNP is omitted if a more significant SNP within 5,000 bp exists. Finally, the set of SNPs that overlapped with HLA loci were removed. After completing this pruning process, we were left with a refined set of 127,061 unique SNPs, representing 4,729 unique diseases/traits. We annotated DNA elements in this study by overlapping them with these unique SNPs. Each DNA element was categorized based on whether it harbored a GWAS SNP.

### GREAT enrichment analysis for conserved human elements

We performed GREAT functional enrichment analysis^49^ via the rGREAT^50^ package on the conserved high CO-V, high CO-B, and gene-homology high CO-V elements in the human genome. For high CO-V or high CO-B elements in the core set, we used the set of all sequence-aligned regions as the background. For gene-homology high CO-V elements in the extended set, we used the set of all DHSs as the background. Due to GREAT’s requirement of a maximum of 600,000 background regions, we subsampled the background regions to this limit. The respective conserved regions within the subsampled background set were then used as the positive set for the analysis.

### Applying FUNCODE to gene expression data

ENCODE RNA-seq data were collected from the ENCODE data portal. All human and mouse polyA+ RNA-seq and total RNA-seq gene quantification data uniformly processed with the ENCODE pipeline were downloaded. For a complete list of the samples included, see **Table S2**. Human and mouse homologous gene annotations (HomoloGene) were downloaded from the Mouse Genome Informatics (www.informatics.jax.org/downloads/reports/HOM_MouseHumanSequence.rpt). Only 16,468 orthologous gene pairs were used. RNA-seq data were integrated using the FUNCODE pipeline with modified parameters. During CCA and the anchor finding step, 3,000 shared variable features were used, and the top 30 canonical variables and five nearest neighbors (k.anchor) were considered. 40 nearest neighbors were used when filtering anchors (k.filter). Log-normalized count data were integrated for visualization in **Fig. 3a**, and the top 30 principal components from the integrated data were used for UMAP visualization. The anchors identified are listed in **Table S1**. Conservation scores for all orthologous gene pairs are available in **Table S5**.

### Associating regulatory elements to their candidate target genes

We employed the activity by contact (ABC) model predictions based on 131 human samples to better associate regulatory elements with their target genes. The ABC model integrates regulatory activity signals and chromatin conformation data to predict gene and regulatory element associations^27,28^. The prediction file (‘AllPredictions.AvgHiC.ABC0.015.minus150.ForABCPaperV3.txt.gz’) was downloaded from https://www.engreitzlab.org/abc/. These predictions integrated tissue or cell-type-specific regulatory element signals with averaged Hi-C loops from 10 human cell types. The dataset included all element-gene connections with ABC scores of 0.015 or higher. Initially, genomic coordinates in the prediction files were converted from the hg19 to the hg38 genome assembly using Liftover. An element from our set is considered mapped to an element in the prediction file if there is an overlap of more than 100 bp. Elements with such mappings are then linked to the corresponding genes in the prediction file.

For mouse regulatory elements, due to the absence of equivalent ABC predictions, we employed a proximity-based approach to link elements to genes. Utilizing the previously defined gene regulatory domains of mouse genes, a mouse element was associated with a gene if it fell within its regulatory domain.

With human and mouse element-gene associations, we then defined consistent target genes for pairs of human-mouse elements. For each pair, we compared whether the list of human and mouse target genes contained any pair of orthologous genes (see earlier section for obtaining the list). If there was at least one orthologous gene pair, the element pair was considered to have at least one consistent target. This distinguished the two types of element pairs in **Fig. 3e** and **Fig. S7a**.

For the pairs with a consistent target, we computed Spearman’s correlation of regulatory element conservation score with the gene expression conservation score (GE-CO-V/B) within different classes of human or mouse cCREs. The result stratified by human cCRE is displayed in **Fig. 3g**, and the one stratified by mouse cCRE is illustrated in **Fig. S7b**.

### Manual matching of human and mouse tissue and cell types

Human and mouse samples were manually matched based on the experiment metadata from the ENCODE data portal (**Table S2**). Two samples were considered a match if their biosample term, life stage (embryonic, child, or adult), and treatment were the same. In total, 20 tissue or cell types were matched for chromatin accessibility, 9 were matched for H3K4me3, 5 were matched for H3K27ac, and 12 were matched for H3K4me1 (**Table S6**). For comparison with the FUNCODE CO-V scores, we computed CO-V scores based on the manually matched samples. A Spearman Correlation was calculated instead of a weighted Spearman Correlation in the manual case.

### Cross-validation of variable and baseline signal in new tissue

To evaluate the conservation scores’ ability to predict conserved variable activities, we conducted a cross-validation study. In each CV partition, the test set comprised two manually matched sample pairs between human and mouse absent from the training set (**Table S9**). FUNCODE conservation scores were computed for the core set of element pairs from the training set data. Each time, 1,000 regulatory element pairs were randomly sampled from the top most conserved pairs, as defined by each conservation score. For evaluation, the fold change between the two held-out samples was first computed for the sampled element pairs of human and mouse separately. Next, we used Spearman’s correlation between human and mouse fold changes to benchmark each score’s ability to predict conserved signals in previously unseen tissue or cell types. This was repeated 20 times and the average correlation was taken. In this comparison, the CO-V scores based on manual matching were also computed with the two holdout samples removed.

To evaluate the conservation scores’ ability to predict conserved baseline activity, one manually matched sample was randomly left out at each fold of the cross-validation (**Table S6**). We then used each conservation score to rank all regulatory element pairs in the core set. Each time, 1,000 regulatory element pairs were randomly sampled from the top most conserved pairs. The top 5% regulatory elements with the highest activities in the core set were taken as highly active elements for the human and mouse samples separately. Among the 1,000 regulatory elements, a Jaccard index was computed, which was the number of elements highly active in both species divided by the number of elements highly active in either species. We used the Jaccard index to benchmark each score’s ability to predict the conservation of baseline regulatory activities. High predictive power in this context implies that if an element is active in one species and conserved, its counterpart should also be active in the other species, regardless of tissue or cell types.

### Evaluation of conservation scores for predicting zebrafish conservations

Similar to human and mouse analysis, we aligned the summits of human DNase I hypersensitive sites (DHSs) to zebrafish genome danRer10 using the pairwise sequence alignment produced by Blastz, downloaded from the UCSC genome browser (https://hgdownload.cse.ucsc.edu/goldenpath/hg38/vsDanRer10/hg38.danRer10.net.axt.gz). Windows of 200 base pairs were created centering on the human DHS summits and the aligned positions in the zebrafish genome. For this analysis, we used the subset of human regulatory elements such that an aligned region exists in both mouse and zebrafish.

All available ATAC-seq and histone ChIP-seq peak files for zebrafish tissues^7^ in the ‘narrowpeaks’ format were downloaded from Gene Expression Omnibus (accession: GSE134055) (https://www.ncbi.nlm.nih.gov/geo/query/acc.cgi?acc=GSE134055). The overlap of the peak regions with the aligned zebrafish regions was computed for each tissue and data modality.

The zebrafish data involved ten tissues: brain, colon, heart, intestine, kidney, liver, muscle, skin, spleen, and testis. They were considered matched to human adult tissue samples with the exact names. For detailed tissue classification of human samples, see **Table S2**. Cell lines and in vitro differentiated samples were not included. For comparing tissue-specific activities, we computed the log2(fold change) of each tissue over all nine other tissues for both human and zebrafish.

For zebrafish, only the presence or absence of peaks were used. For evaluation, the regions that can be aligned to both zebrafish and mouse were first filtered by different conservation scores. Each time, 1,000 random regions were sampled from the set that passed the filter, for which Spearman’s correlation of the human and zebrafish fold changes were computed.

For comparing tissue invariant baseline activities, we repeated the same filtering step by conservation scores and evaluated by the fraction of zebrafish samples (out of ten tissues) that had a peak overlapping the region.

### Transfer of ENCODE Hi-C interactions between human and mouse

Long-range chromatin interactions or loops (in .bedpe format) of 17 human and 9 mouse experiments were downloaded from the ENCODE data portal (**Table S2**). Among the loops identified in the human Hi-C experiments, a subset featured high-resolution loop anchors with widths significantly narrower than the low-resolution anchors (500 bp vs 10,000 bp). During the transfer of human loops to mouse, we focused on the loops with high-resolution anchors to enable more precise association between regulatory elements and loop anchors. For the mouse data, the loop anchors were broader due to the use of an older protocol^35^ [ENCODE tracking # ENC4P10]. To ensure the accuracy of associations with regulatory elements, we concentrated on loops whose anchors overlapped no more than five regulatory elements.

The transfer pipeline involved three main steps. First, we mapped regulatory elements to loop anchors and established interactions between these elements. An interaction between a pair of regulatory elements was inferred if a Hi-C loop existed with two anchors coinciding with each of the two elements. Next, we mapped the regulatory elements using regulatory element pairs (REPs). The mapping was determined by the REPs associated with each element within the core or extended set. Elements in the core set were mapped to their sequence-aligned counterparts and further stratified based on the presence or absence of FUNCODE conservation. Elements in the extended set, not included in the core set, could be associated with multiple REPs, which might or might not share transcription factor binding site (TFBS) motifs. In such cases, REPs with shared TFBS motifs were prioritized. We further annotated each transferred interaction by the presence or absence of CTCF binding motifs, focusing on loops where at least one anchor contained a CTCF binding motif. We removed all transferred interactions whose anchors were on different chromosomes.

Finally, we assessed the transferred interactions by checking for the existence of a loop in the corresponding cell type of the other species that connected them. Evaluations were conducted for experiments on CD4^+^ T cells and B cells, as these cell types were available for both species. Differences in Hi-C protocols led to significant differences in the distance between loop anchors between the human and mouse Hi-C data. Specifically, only 7% of interactions identified from mouse Hi-C had anchor distances smaller than 65,000 bp, compared to 41% in the human-transferred interactions. Thus, we restricted our evaluation to interactions with anchor distances greater than 65,000 bp to ensure a fair comparison. Transferred loops below this threshold might still be accurate but were unlikely to be detected by the mouse Hi-C experiments. Similarly, 0.26% of loops in human Hi-C data had distances greater than 4,000,000 bp compared to 16% in mouse-transferred interactions, prompting us to limit evaluations to interactions with distances smaller than 4,000,000 bp. We summarized the average fraction of validated interactions for each combination of the REPs categories used to transfer the loop (**Fig. 5d, S9**). Overall, transfers where both anchors were transferred by sequence-aligned and FUNCODE conserved REPs demonstrated the highest validation rate. We transferred loops from 17 human experiments covering eight immune cell types to the mouse genome and reported the sequence-aligned and FUNCODE conserved interactions as the FUNCODE-predicted chromatin interactions (**Fig. 5e**, **Table S8**).

### Single-cell ATAC-seq data processing and integration

We downloaded BAM files for mouse sciATAC from the Mouse ATAC Atlas^36^ (https://atlas.gs.washington.edu/mouse-atac/data/) and fragment files for human sciATAC from the Human Enhancer Atlas^37^ (http://catlas.org/humanenhancer/). The reads were first processed by counting the number of reads for pairs of regulatory elements in our core set (sequence-aligned DNA elements). Mouse sciATAC-seq reads were originally aligned to the mm9 genome. Since mouse regulatory elements in our core set are defined on the mm10 genome, we first converted the mm10 regulatory elements to mm9 using liftOver. From these atlases, we identified four common tissues: lung, liver, heart, and large intestine. The specific samples included were:

- <u>Human</u>: lung (SM-A62E9, SM-A8WNH), liver (SM-A8WNZ), heart (lv_SM-IOBHO, lv_SM-JF1NY, atrial appendage_SM-JF1NX, atrial appendage_SM-IOBHN), colon (transverse_SM-A9HOW, transverse_SM-A9VP4, transverse_SM-ACCQ1, transverse_SM-BZ2ZS, transverse_SM-CSSDA, sigmoid_SM-AZPYO, sigmoid_SM-JF1O8).
- <u>Mouse</u>: Lung1_62216, Lung2_62216, Liver_62016, HeartA_62816, LargeIntestineA_62816, LargeIntestineB_62816.

Human and mouse cell types, as labeled in the original publications, were mapped according to **Table S10.**

We binarized the signal by assigning a value of one to all peaks with more than one read and excluded cells with fewer than 900 non-zero elements. We retained only cell types with matched labels across species, comprising 77.3% of human cells and 79.8% of mouse cells. To select the initial feature set for data integration, we ranked each regulatory element based on the count of cells with non-zero reads for the element, doing this separately for human and mouse. The human and mouse ranks were then added, and the top 300,000 elements with the highest rank sums were selected. In this set, each regulatory element was detected in a median of 100 human cells (0.14%) and 99 mouse cells (0.46%). For the control method, we further refined our feature selection using the same rank sum, narrowing it down to the top 25,000 or 48,160 features. These features represent the regulatory elements most commonly shared between the human and mouse datasets but not necessarily the most conserved in terms of context-dependent activities. To apply conservation scores, the initial set of features was further ranked based on each score, and the top 25,000 or 48,160 features were chosen correspondingly.

The integration was performed using the default Seurat pipeline. We considered the top 30 canonical variables and 40 nearest neighbors (k.anchor) during Canonical Correlation Analysis (CCA). For anchor filtering (k.filter), 100 nearest neighbors were considered. Subsequently, the ‘IntegrateData’ function was used with the top 30 canonical variables and 100 neighbors (k.weight) for weighting the anchors. Following data integration, we applied latent semantic indexing (LSI) to reduce the dataset to 30 dimensions and used UMAP for visualization.

We used a label transfer procedure based on mutual nearest neighbors (MNN) to evaluate integration accuracy. Specifically, we treated human and mouse datasets as distinct in the integrated LSI space and identified their MNNs, considering up to 40 nearest neighbors for each. The labels of mouse cells were computed as Gaussian kernel weighted averages of the human anchors’ labels, where the Gaussian kernel function (with a standard deviation of 20) took distances to anchors as input and considered up to 100 mutual neighbors. We calculated the label prediction matrix by computing the weighted average of the label indicators of the anchors, with the transferred label being the one with the highest predicted probability.

We then computed differential signals between two cell types for all elements detected in at least one cell in both species. We first grouped single cells into pseudo-bulk and computed the average signals by summing up counts and log2-normalizing the count vector. The differential signal vector was then calculated as the difference between the two pseudo-bulk vectors for the cell types. True conserved differential elements were identified as those with both human and mouse absolute differential signals greater than 3 (an eightfold change) and of the same sign. For evaluation, we ranked the elements based on human and mouse differential signals separately and then added the ranks to obtain the overall rank sum. Finally, we computed the fraction of true conserved elements among the top-ranked elements for different rank sum cutoffs.

## Data Availability

Conservation scores are available via the ENCODE data portal (https://www.encodeproject.org/search/?type=Annotation&annotation_type=cross-species+functional+conservation), and listed in **Table S3**. Score track hubs for the UCSC genome browser are available on GitHub (https://github.com/wefang/funcode). FUNCODE scores for regulatory elements are accessible by query through the web application (https://jhubiostatistics.shinyapps.io/FUNCODE).

## Code Availability

The code for the FUNCODE pipeline and scripts to reproduce the analysis are on (https://github.com/wefang/funcode).

## Author Contributions

H.J. conceived the study. W.F. conducted all the analyses. C.C. compiled the core set of element pairs and existing conservation scores. C.C. also processed the Activitiy by Contact data. B.Z. compiled the results for the extended set of regulatory element pairs. R.Z. helped with single-cell integration. W.Z. and Y.W. mapped TFBS motifs to human and mouse genomes. W.F. and H.J. wrote the initial manuscript. C.C., B.Z., R.Z., Y.W. and W.Z. reviewed the manuscript and provided feedback. All authors approved the final manuscript.

## Acknowledgments

We thank Drs. Timothy E Reddy, Ali Mortazavi, Anshul Kundaje, Erez Lieberman Aiden, Michael Pazin, Kushal Kumar Dey, Jill Moore, and the ENCODE mouse working group for their helpful discussions and feedback. We also thank Drs. Benjamin Hitz, Ingrid Ashley Youngworth and Idan Gabdank from the ENCODE Data Coordination Center (DCC) for assisting with the FUNCODE data release via ENCODE data portal. This work was supported by the National Institutes of Health (NIH) grants R01 HG010889, R01 HG009518 and R01 HG013409. We acknowledge the NIH/NHGRI for their crucial support in advancing this research. The computational work for this project was conducted using the resources of the Johns Hopkins Joint High-Performance Computing Exchange (JHPCE) and the Maryland Advanced Research Computing Center (MARCC).

## Supplementary figures

**Fig. S1.**
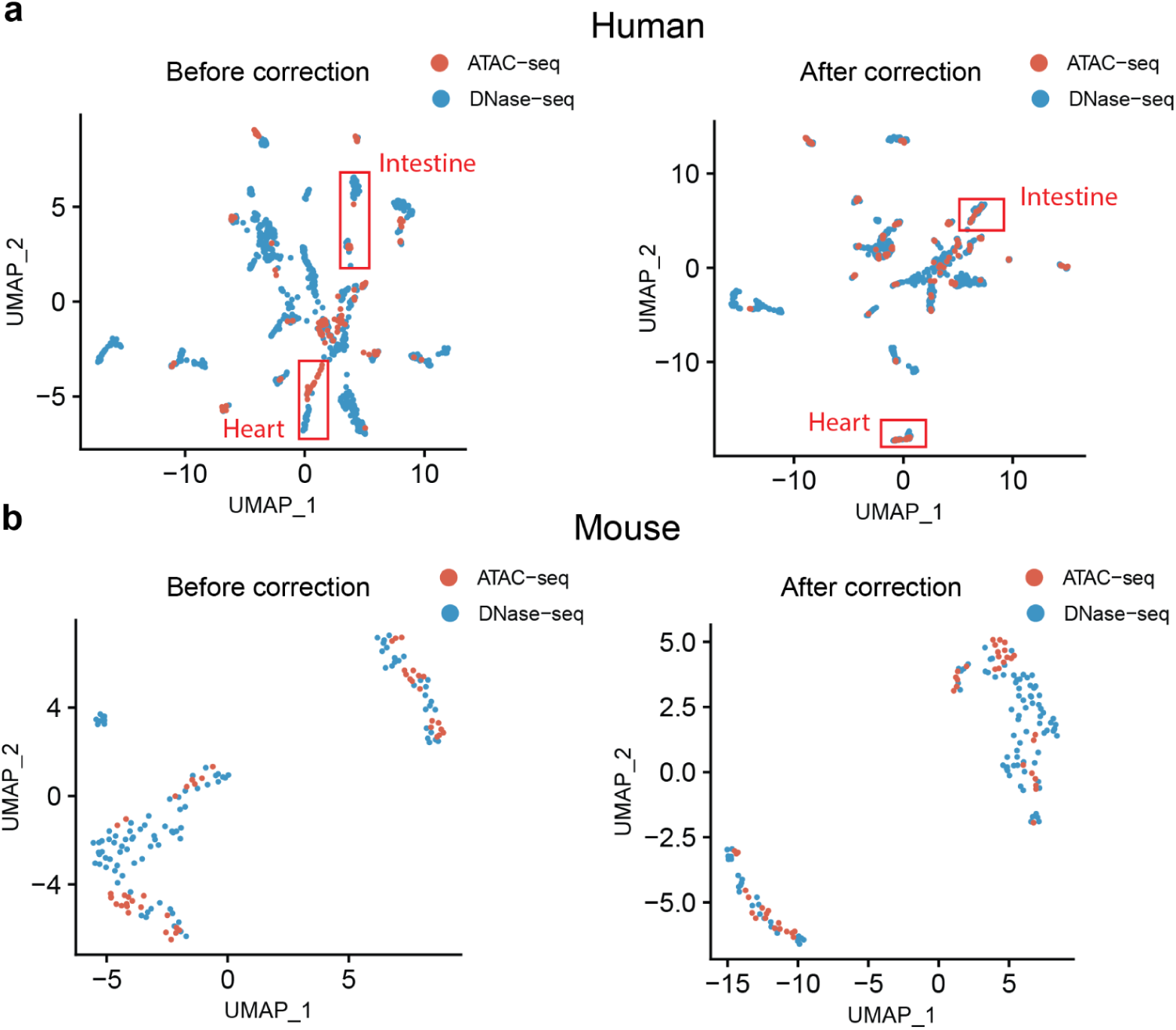
Correcting assay-specific effects between DNase-seq and ATAC-seq samples. (**a**) Scatterplots showing UMAP embeddings of human DNase-seq and ATAC-seq samples before and after assay-specific effect correction. Heart and intestine tissues are marked in red boxes as examples. Samples clustered more according to tissue than to assay after correction. (**b**) Same as (**a**), but for mouse samples. No significant difference in cell type clustering was observed before and after correction.

**Fig. S2.**
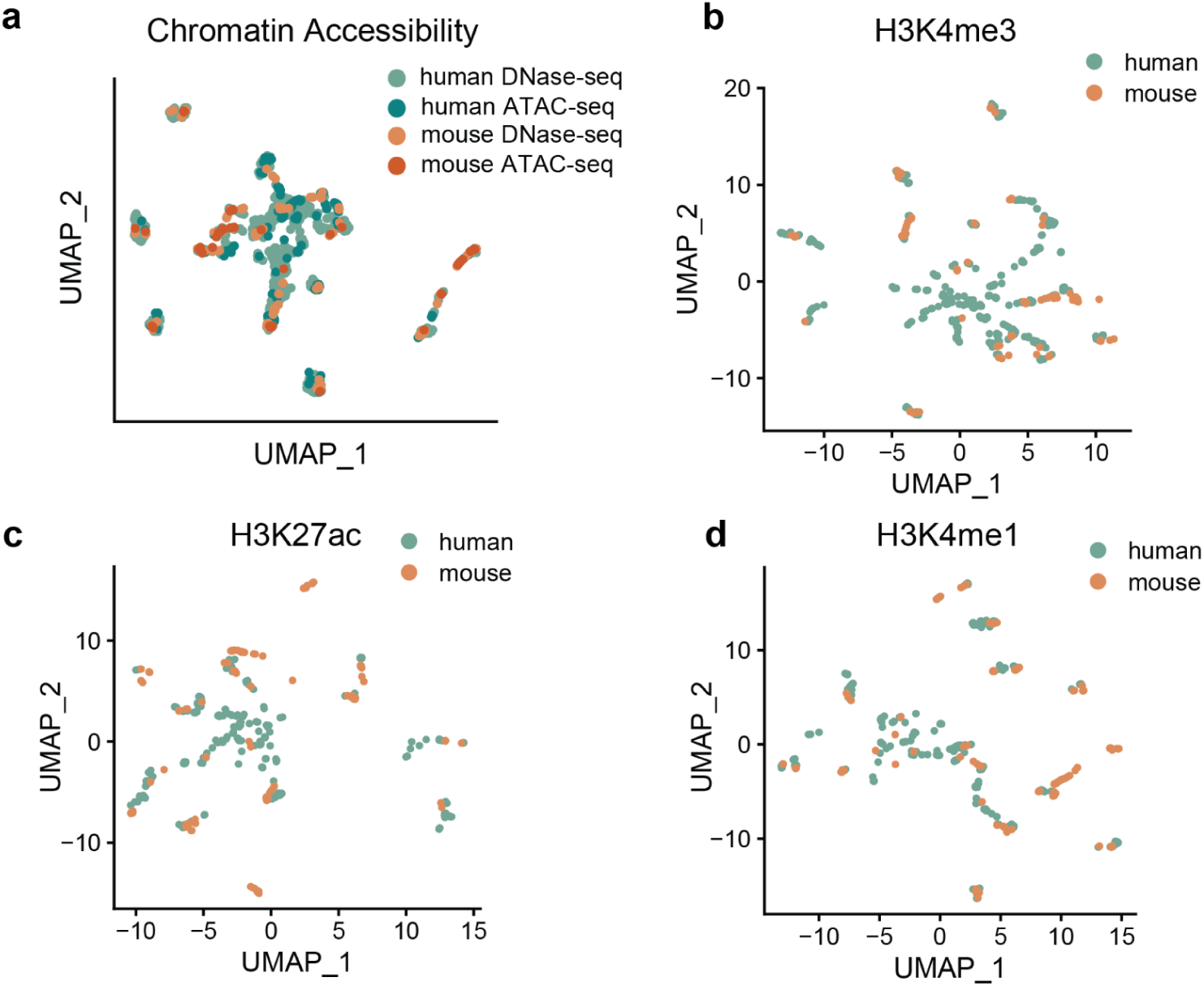
Integration of human and mouse samples from ENCODE for four data modalities. (**a**) Scatterplot showing UMAP embeddings of human and mouse chromatin accessibility (DNase-seq and ATAC-seq) data. (**b**) Same as (**a**), but with H3K4me3 Histone ChIP-seq data. (**c**) Same as (**a**), but with H3K27ac Histone ChIP-seq data. (**d**) Same as (**a**), but with H3K4me1 Histone ChIP-seq data. For i*n silico* matched human and mouse samples of each data modality, see **Table S1**.

**Fig. S3.**
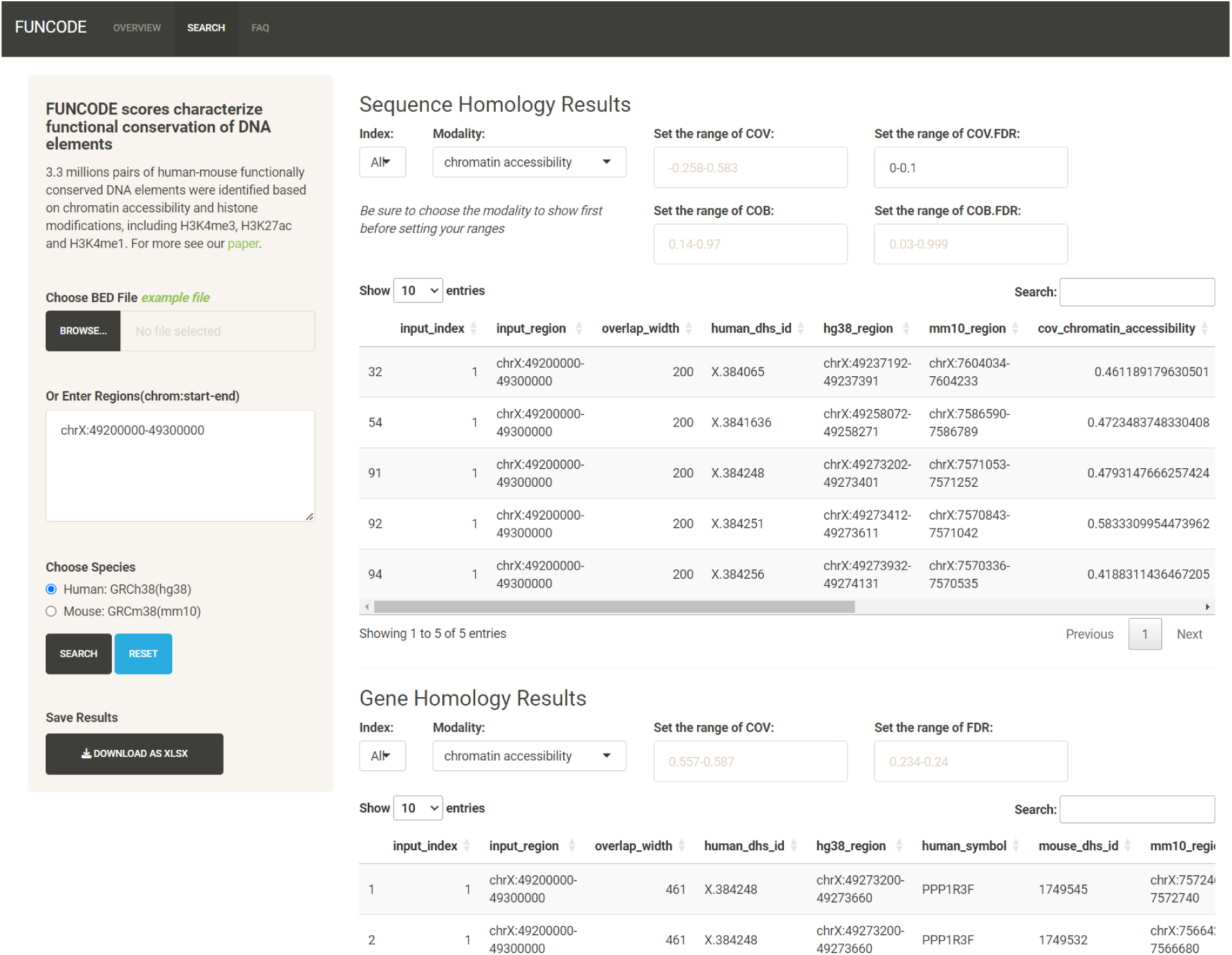
FUNCODE web interface. Screenshot displaying the FUNCODE web application interface, enabling users to search for conserved regulatory element pairs (REPs) in human or mouse genomes. Results are presented in tables, with options to filter by data modality, score, or false discovery rate.

**Fig. S4.**
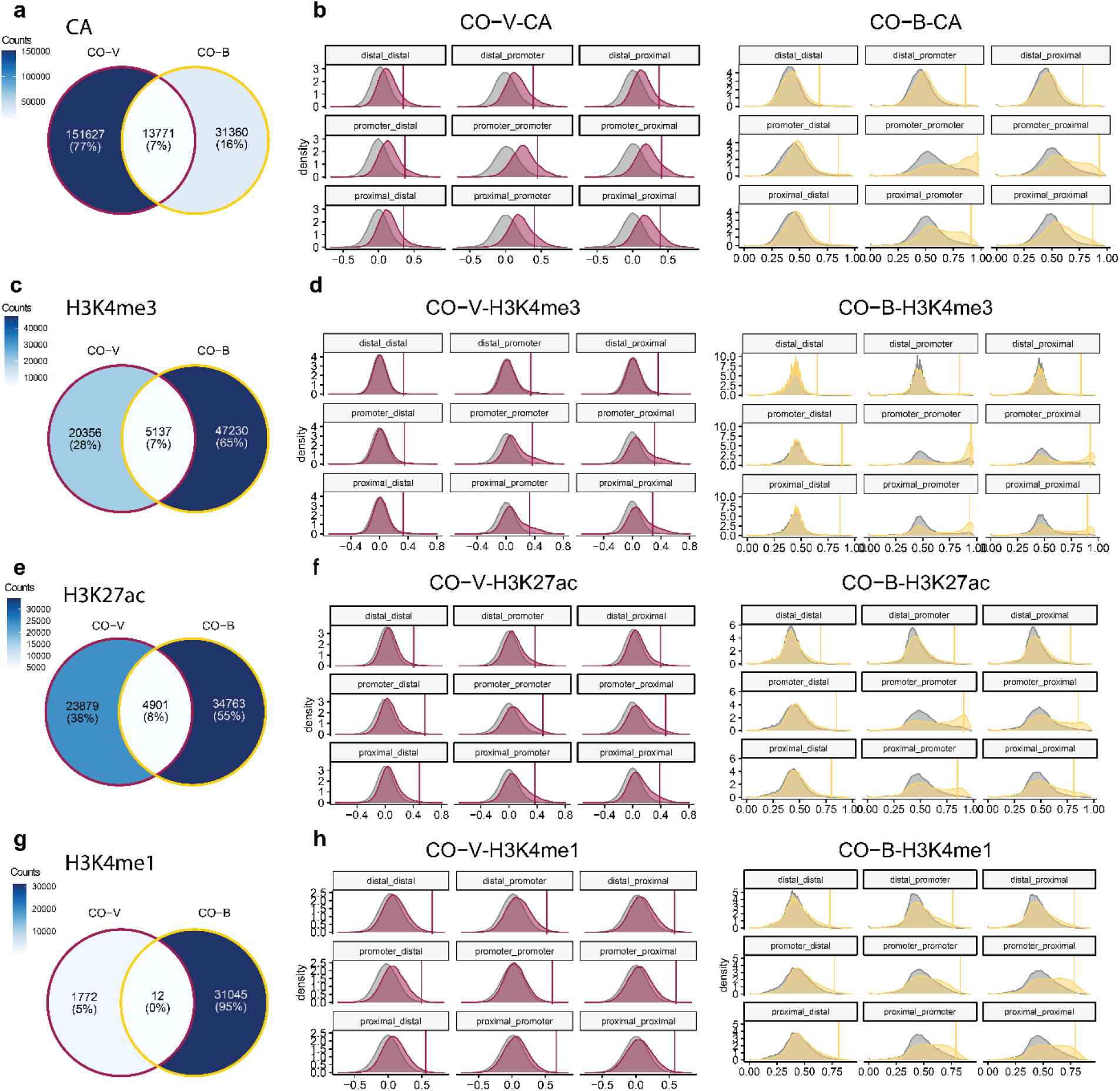
Calling significantly high CO-V and CO-B elements for sequence homology element pairs (core set). (**a**) Venn diagrams showing the amount of overlap between sequence homology CO-V-CA and CO-B-CA elements. (**b**) Density plots showing the distribution of CO-V-CA scores (left) and CO-B-CA (right) in nine gene context pair categories. For example, the ‘promoter-proximal’ category includes pairs of DNA elements where the human element is a promoter and the mouse element is a proximal enhancer. The null distributions in each category (created by random pairing of human and mouse elements with sequence homology) are depicted in gray. Vertical lines indicate the significance cutoff at a 10% false discovery rate (FDR). (**c**,**e**,**g**) Same as (**a**), but for H3K4me3, H3K27ac, and H3K4me1 histone modification conservation, respectively. (**d**,**f**,**h**) Same as (**b**), but for H3K4me3, H3K27ac, and H3K4me1 histone modification conservation, respectively.

**Fig. S5.**
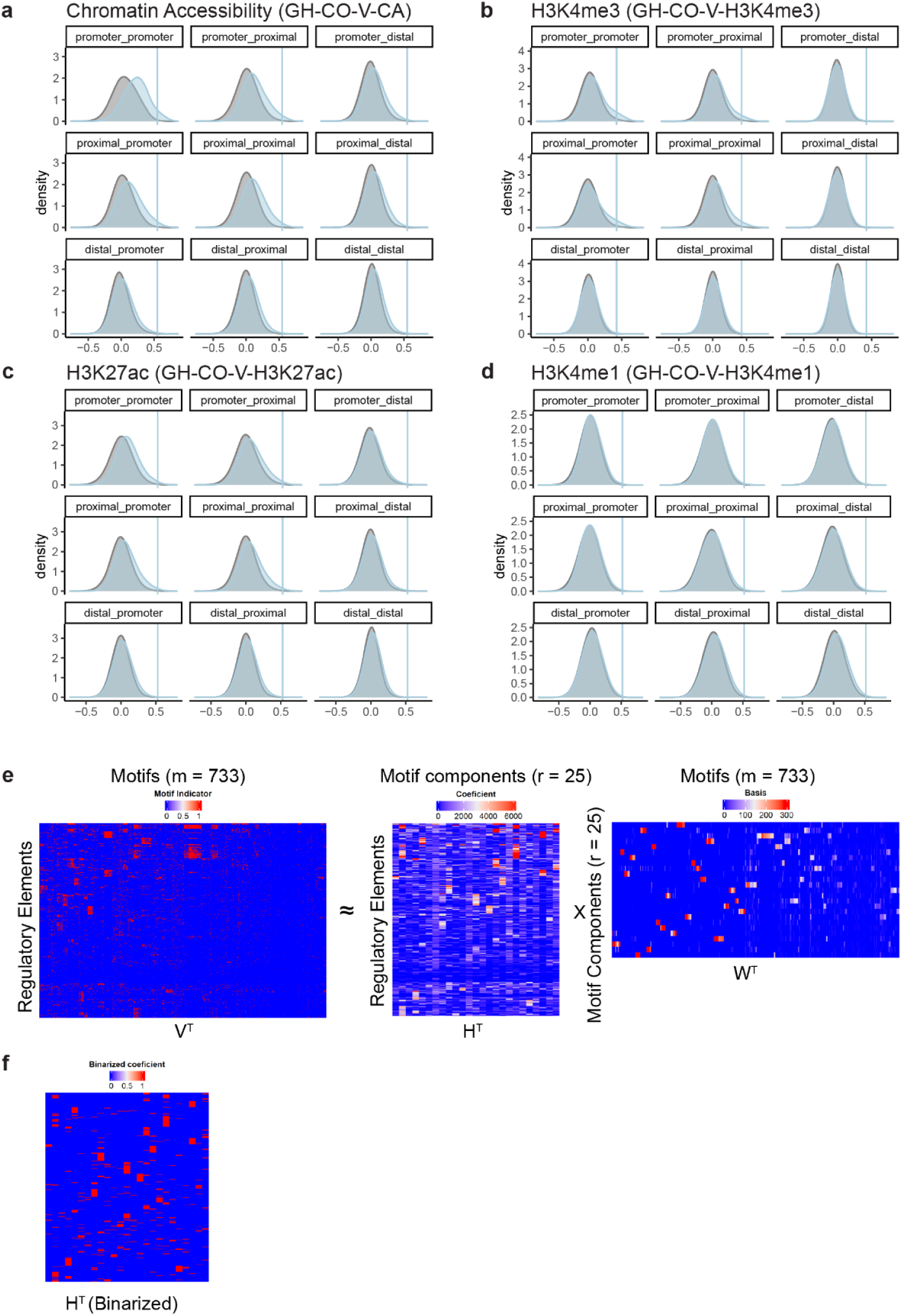
Calling significantly high CO-V elements for gene homology element pairs (extended set) and annotating shared transcription factor binding site motifs. (**a**) Density plots showing the distribution of GH-CO-V-CA scores in nine gene context pair categories. For example, the ‘promoter-proximal’ category includes pairs of DNA elements where the human element is a promoter and the mouse element is a proximal enhancer. The null distributions in each category (created by random pairing of human and mouse elements with gene homology) are depicted in gray. Vertical lines indicate the significance cutoff at a 25% false discovery rate (FDR). (**b**,**c**,**d**) Same as (**a**), but for H3K4me3, H3K27ac, and H3K4me1 histone modification conservation, respectively. (**e**) Heatmaps illustrating reducing the transcription factor indicator matrix (*V*^*T*^) by non-negative matrix factorization (NMF). The indicator matrix is a binary matrix indicating if a TFBS motif is mapped to a regulatory element. It is expressed as the matrix product of the coefficient matrix (*H*^*T*^) and the basis matrix (*W*^*T*^). (**f**) Heatmap displaying a binarized coefficient matrix indicating the association between regulatory elements and TFBS motif components. *HH*^*TT*^ is binarized by taking a row-wise z-score cutoff of 2.0.

**Fig. S6.**
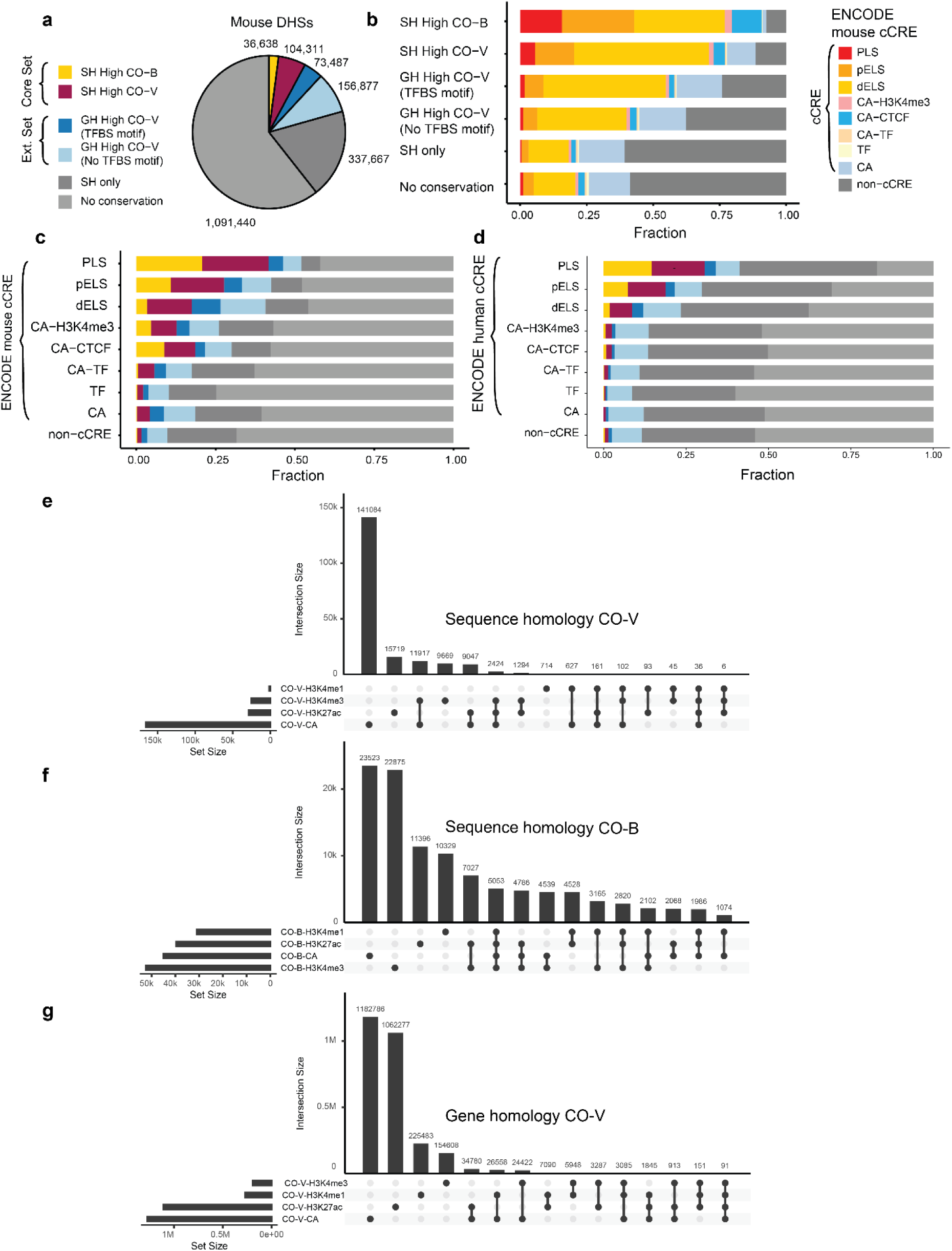
Characteristics of conserved elements and overlaps of CO-V and CO-B elements called using different data modalities. (**a**) Pie chart showing the fraction of different conserved elements among all mouse DHSs. Color indicates four different conservation categories. Pairs that are both high CO-V and high CO-B are shown as high CO-V. For the amount of overlap, see **Fig. S4**. (**b**) Stacked barplot showing the enrichment of mouse candidate Cis-Regulatory Elements (cCRE) in different conservation categories. (**c**) Stacked barplot showing the enrichment of conserved elements in different mouse cCRE classes. Element pairs conserved with both sequence homology and gene homology were shown as sequence homology. See panel (**a**) for color legend. (**d**) Stacked barplot showing the enrichment of conserved elements in different human cCRE classes. Element pairs conserved with both sequence homology and gene homology were shown as sequence homology. See panel (**a**) for color legend. (**e**) Upset plot showing the amount of intersection of high CO-V elements of four different data modalities (chromatin accessibility, H3K4me3, H3K27ac, and H3K4me1) in the core set (sequence homology). (**f**) The same as (**e**), but for high CO-B elements in the core set (sequence homology). (**g**) The same as (**e**), but for high CO-V elements in the extended set (gene homology).

**Fig. S7.**
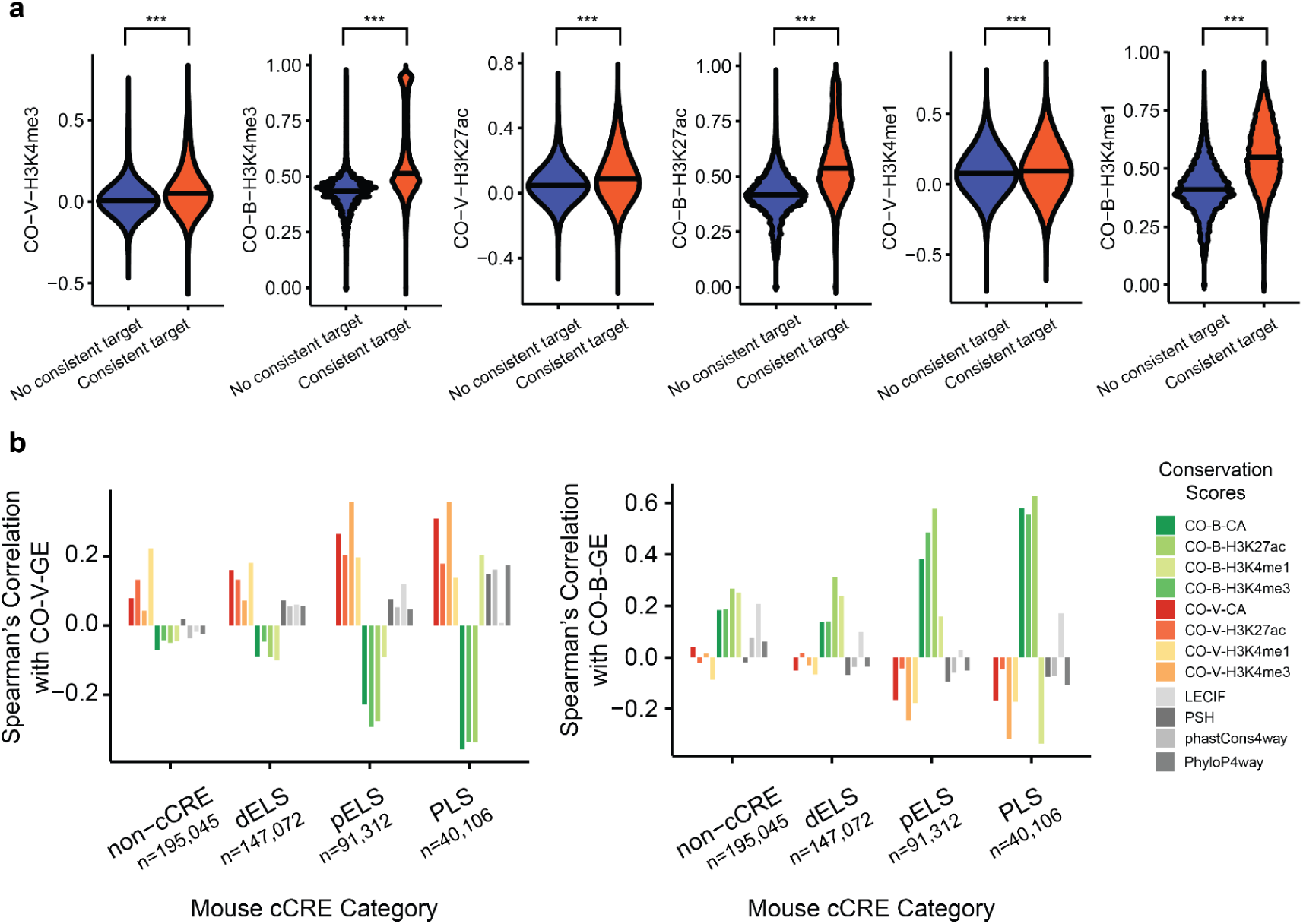
Regulatory element conservation scores of different modalities predict gene expression conservation scores. (**a**) Violin plots showing the distributions of different FUNCODE scores. X-axis and color indicate pairs with or without consistent target genes. Pairs with a consistent target gene had higher FUNCODE scores than pairs without. ***: Wilcoxon rank sum test, *p* < 0.001. (**b**) Barplots showing the Spearman’s correlation (y-axis) between different regulatory element conservation scores and gene expression conservation scores (left: CO-V-GE, right: CO-B-GE) of the consistent target gene pairs across different classes of mouse cCRE (x-axis). CO-V scores of regulatory elements showed the highest correlation with CO-V-GE and an inverse correlation with CO-B-GE. CO-B scores of regulatory elements were most correlated with CO-B-GE and inversely correlated with CO-V-GE. The strength of these correlations was higher in PLS (promoter), followed by pELS (proximal enhancer), dELS (distal enhancer), and lowest in non-cCRE.

**Fig. S8.**
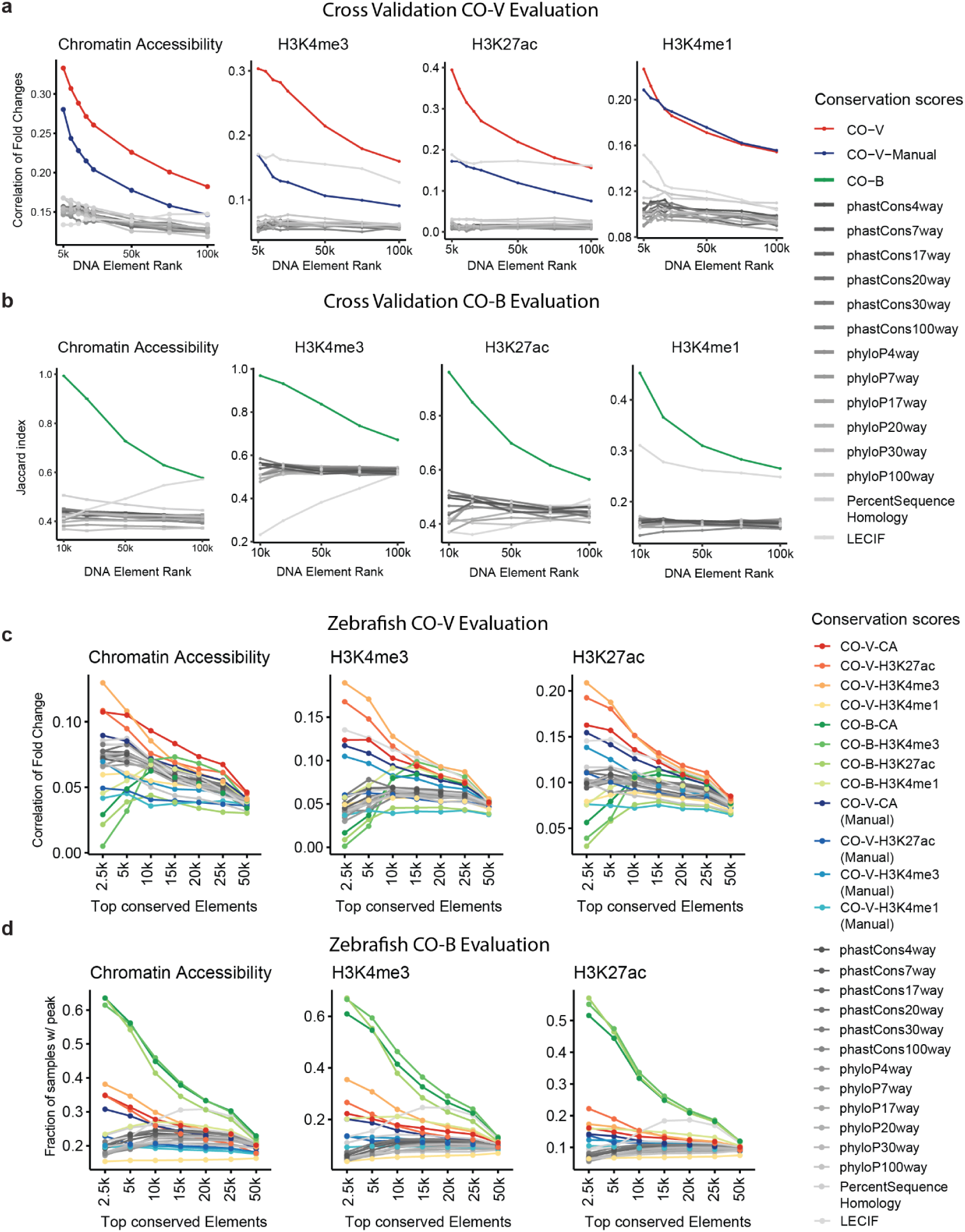
Extended evaluations of FUNCODE scores for predicting conserved activities in new tissue, cell types, or species, including additional data modalities and sequence-based conservation scores. (**a**) Line plots showing conservation of differential activities between new tissue or cell types (as measured by correlation of human and mouse fold changes, y-axis) at different conservation score cutoffs (x-axis). The CO-V score in each panel is the CO-V score specific to the data modality. (**b**) Line plots showing conservation of baseline activities in new tissue or cell types (as measured by the Jaccard index on the y-axis) at different conservation score cutoffs (x-axis). The CO-B score in each panel is the CO-B score specific to the data modality. (**c**) Line plots showing conservation of tissue-specific activities between human and zebrafish (as measured by correlation of human and zebrafish fold changes, y-axis) at different human-mouse conservation score cutoffs (x-axis). (**d**) Line plots showing conservation of baseline activities (as measured by the fraction of zebrafish samples with an overlapping peak, y-axis) at different human-mouse conservation score cutoffs (x-axis).

**Fig. S9.**
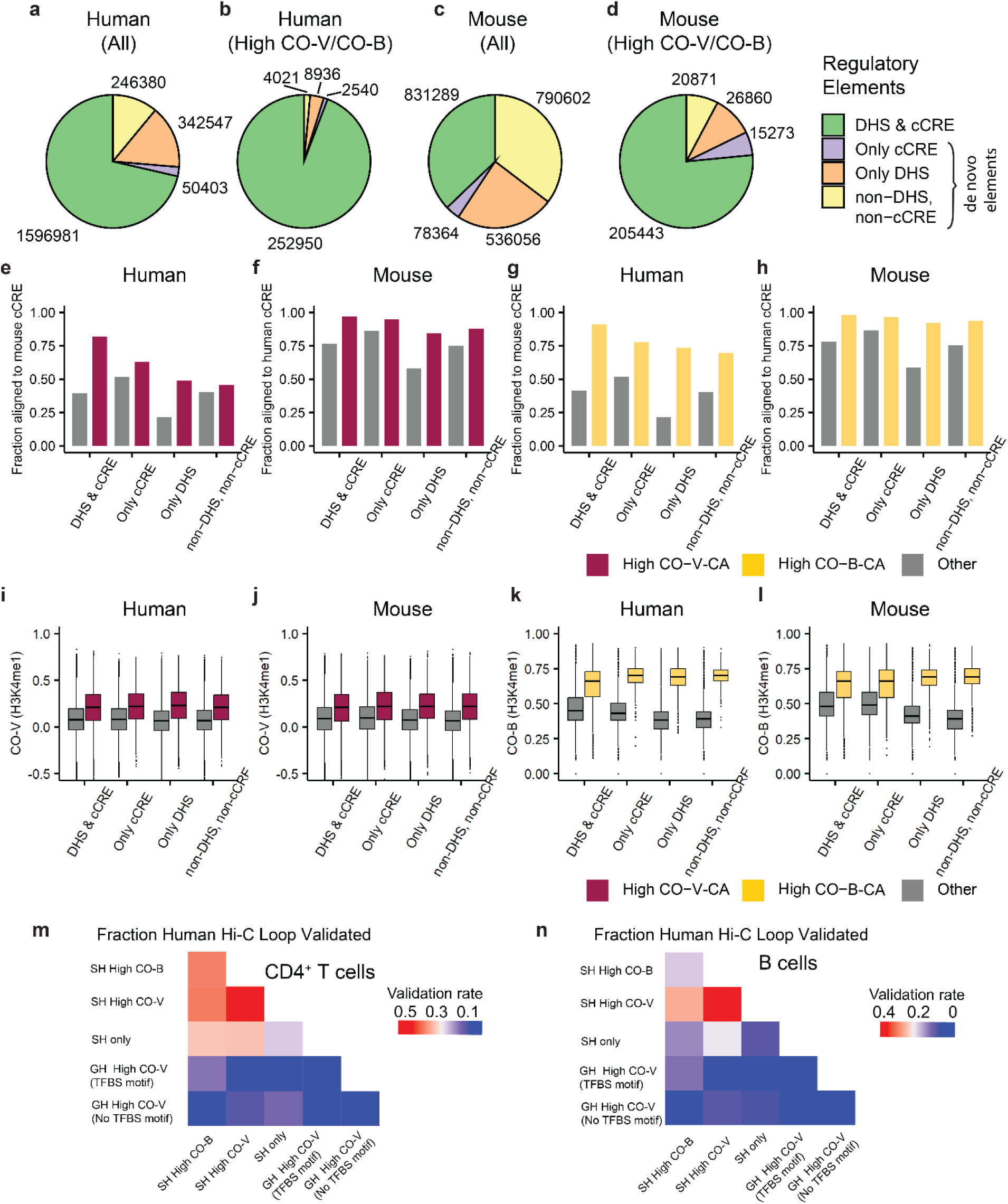
Using FUNCODE for de novo functional annotations in the human and mouse genome. (**a**) Pie chart breaking down current human functional annotations for all elements in the core set. (**b**) Pie chart breaking down current human functional annotations for FUNCODE conserved elements in the core set. A larger fraction of conserved elements were annotated. However, a significant portion (non-green colors) was unannotated and are likely functional elements. (**c**) Pie chart breaking down current mouse functional annotations for all elements in the core set. (**d**) Pie chart breaking down current mouse functional annotations for FUNCODE conserved elements in the core set. A similar trend as in (**b**) is observed for mouse. (**e**) Barplot showing alignment rate of human DNA elements to mouse cCRE (y-axis) for different categories in (**a**) or (**b**) (x-axis). Color indicates high CO-V elements versus others. Conserved (high CO-V) but unannotated elements showed comparable rates of alignment to functional elements compared to conserved and annotated elements. Both were higher compared to non-conserved elements, irrespective of being annotated or not. (**f**) Barplot showing alignment rate of mouse DNA elements to human cCRE (y-axis) for different categories in (**c**) or (**d**) (x-axis). Color indicates high CO-V elements versus others. (**g**) Same as (**e**), but color indicates high CO-B elements versus others. (**h**) Same as (**f**), but color indicates high CO-B elements versus others. (**i**) Boxplot showing the distribution of CO-V-H3K4me1 scores (y-axis) for different categories of human functional annotations (x-axis). Color indicates high CO-V elements versus others. Conserved (high CO-V, defined without using H3K4me1 ChIP-seq data) but unannotated elements had similar CO-V-H3K4me1 scores compared to conserved and annotated elements. Both were higher compared to non-conserved elements, irrespective of being annotated or not. (**j**) Same as (**i**), but the x-axis breaks down mouse functional annotations. (**k**) Same as (**i**), but color indicates high CO-B elements versus others. (**l**) Same as (**i**), but the x-axis breaks down mouse functional annotations, and color indicates high CO-B elements versus others. (**m**) Heatmaps showing the validation rates of mouse chromatin interactions transferred from human ENCODE Hi-C data in CD4^+^ T cells. Each cell represents a unique combination of REPs for mapping the loop anchors across species. (**n**) Same as (**m**), but for B cells.

**Fig. S10.**
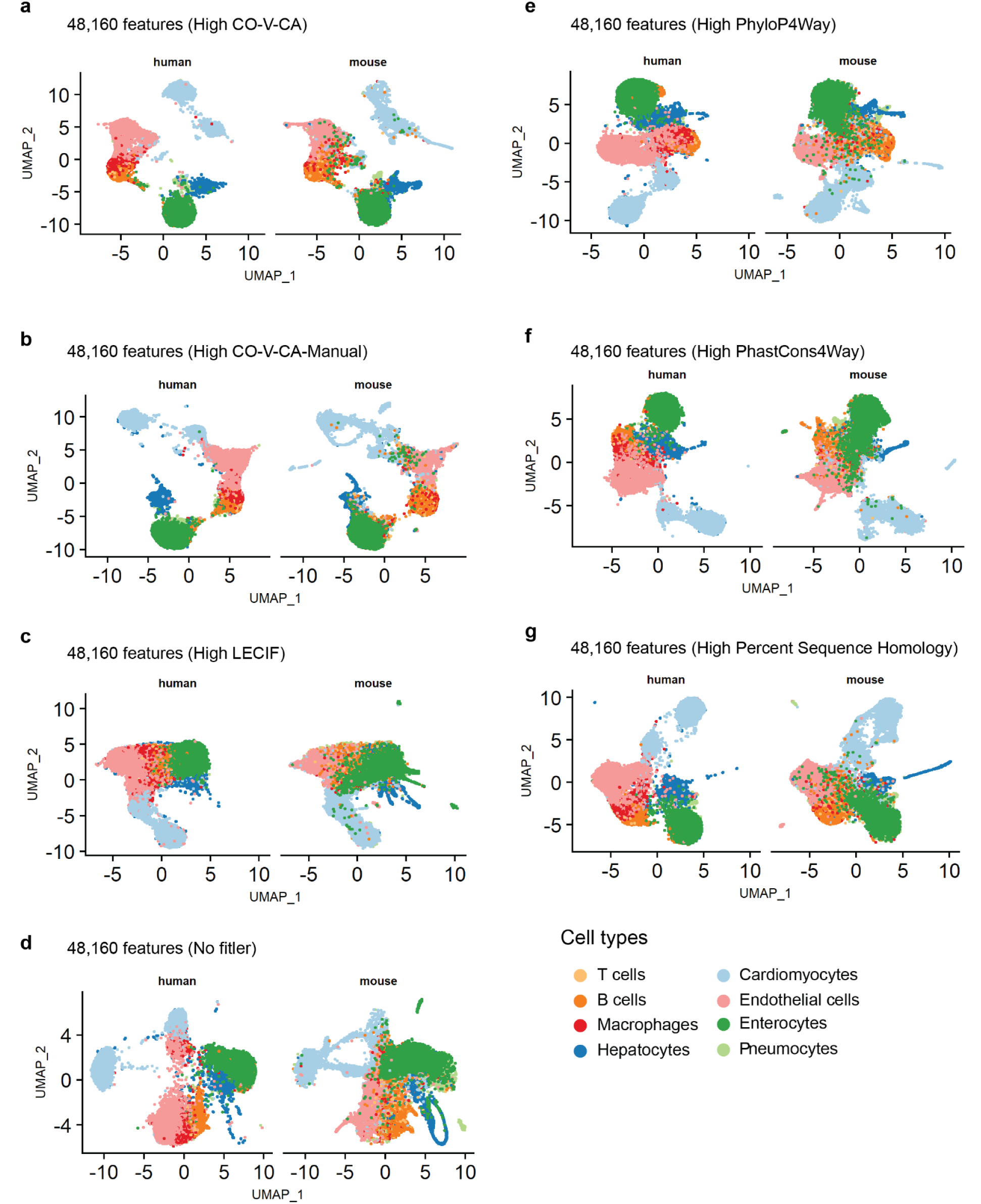
Integration of human and mouse sciATAC-seq data using features filtered by different conservation scores. (**a**) Scatterplot showing embedded single cells from human and mouse sciATAC-seq datasets after cross-species integration. Human and mouse cells were shown in separate panels. Color indicates cell types labeled in the original publications. 48,160 features (sequence-aligned element pairs) with the highest CO-V-CA scores were used for the integration. (**b**) Same as (**a**), but using features with highest CO-V-CA-Manual scores. (**c**) Same as (**a**), but using features with highest LECIF scores. (**d**) Same as (**a**), but no filtering of conservation scores was applied. (**e**) Same as (**a**), but using features with the highest PhyloP4Way scores. (**f**) Same as (**a**), but using features with the highest PhastCons4Way scores. (**g**) Same as (**a**), but using features with highest percent sequence homology (PSH) scores.

## List of supplementary tables

**Table S1**. *In silico* matched human and mouse samples for DNase-seq, ATAC-seq, Histone ChIP-seq, and RNA-seq samples.

**Table S2**. Experimental metadata for ENCODE DNase-seq, ATAC-seq, Histone ChIP-seq, RNA-seq, ENCODE Hi-C, and in situ Hi-C samples.

**Table S3**. FUNCODE score files on ENCODE data portal.

**Table S4**. Enriched GO terms for conserved regulatory elements from GREAT analysis.

**Table S5**. FUNCODE conservation scores for ortholog gene pairs.

**Table S6**. Manually matched samples for DNase-seq, ATAC-seq, and Histone ChIP-seq.

**Table S7**. Conserved and unannotated regulatory elements.

**Table S8**. Transferred ENCODE Hi-C contact loops in the mouse genome.

**Table S9**. Held-out samples in cross-validation analysis.

**Table S10**. Matched cell type labels between human and mouse single-cell ATAC-seq data.

